# An endogenous retroviral element co-opts an upstream regulatory sequence to achieve somatic expression and mobility

**DOI:** 10.1101/2025.01.02.631056

**Authors:** Natalia Rubanova, Darshika Singh, Louis Barolle, Fabienne Chalvet, Sophie Netter, Mickaël Poidevin, Nicolas Servant, Allison J. Bardin, Katarzyna Siudeja

**Affiliations:** Institut Curie, PSL Research University, CNRS UMR 3215, INSERM U934, Stem Cells and Tissue Homeostasis Group, 75005 Paris, France; Institut Curie Bioinformatics core facility, PSL Research University, INSERM U900, MINES ParisTech, 75005 Paris, France; Institute for Integrative Biology of the Cell (I2BC), INSERM U1280, CEA, CNRS, Université Paris-Saclay, Gif-sur-Yvette, France; Department of Biology, University of Versailles St-Quentin, Versailles, France

**Author notes:** These authors contributed equally (Darshika Singh, Louis Barolle).

## Abstract

Retrotransposons, multi-copy sequences that propagate *via* copy-and-paste mechanisms involving an RNA intermediate, occupy large portions of all eukaryotic genomes. A great majority of their manifold copies remain silenced in somatic cells, nevertheless, some are transcribed, often in a tissue specific manner, and a small fraction retains its ability to mobilize. Retrotransposon expression or mobility are increasingly recognized to contribute to normal development and tissue homeostasis, as well as to aging and disease. While it is well characterized that retrotransposon sequences may provide *cis* regulatory elements for neighboring genes, how their own expression and mobility are achieved in different somatic contexts is not well understood. Here, using long-read DNA sequencing, we characterize somatic retrotransposition in the *Drosophila* intestine. We show that retroelement mobility does not change significantly upon aging and is limited to very few active sub-families of retrotransposons. Importantly, we identify a polymorphic donor locus of an endogenous LTR retroviral element *rover*, active in the intestinal tissue. We reveal that gut activity of the *rover* donor copy depends on its genomic environment. Without affecting local gene expression, the copy co-opts its upstream enhancer sequence, rich in transcription factor binding sites, for somatic expression. Further we show that *escargot,* a snail-type transcription factor critical for gut progenitor cell function, can drive transcriptional activity of the active *rover* copy. These data provide new insights into how locus-specific features allow active retrotransposons to produce functional transcripts and mobilize in a somatic lineage.

## Introduction

Transposable elements (TEs), repetitive genetic elements capable of self-replicating and moving from one position in the genome to another, represent a large part of eukaryotic genomes (1). Class I long terminal repeat (LTR) and long interspersed nuclear element-1 (L1) retroelements mobilize by a copy-and-paste mechanism through an RNA intermediate. They are often present in thousands of copies, following what are believed to be waves of mobilization in genomes that date back millions of years (e.g HERV-K elements in human (2)). Many copies subsequently lose DNA sequence integrity over time by acquiring mutations that inactivate their mobilization machinery or are eliminated from genomes through recombination events. Thus, only a small number of copies remain intact and transposition-competent. Additionally, to protect genome integrity, host organisms have developed multiple pre- and post-transcriptional mechanisms of repression operating in the germline and somatic tissues (3,4).

The impact of TEs on evolution and their vast contribution to genome regulation and cell biology is now widely recognized. TE sequences can be repurposed for the benefit of the host, can introduce heritable mutations that cause diseases, contribute to species variation and regulate cellular transcriptional programs (e.g. development, neural progenitor cells) (reviewed in 2). In addition, through the somatic mobility, they are a source of genetic mosaicism that may facilitate tumorigenesis and is of yet unclear role in healthy tissues. Moreover, apart from their mobility, TE RNA or protein products are believed to contribute to important biological processes, such as normal development (5,6) or immune response (7–9). Indeed, TE transcripts, including those with intact ORFs, are repeatedly detected in many tissue types, often with tissue- or cell-type specific expression patterns (10–19). Nevertheless, how TE expression and mobility are regulated in diverse somatic context is not well understood.

Addressing TE regulation in the soma is complex due the highly repetitive nature of TE sequences, which complicates bioinformatic analysis (20,21). Indeed, TE transcripts are derived only from a small subset of active, derepressed loci (16–18,22–26). Additionally, a further challenge is the difficulty in detecting somatic TE mobility, which occurs in small proportion of cells and thus represents only rare events. To date, studies on somatically active TE loci have largely focused on mammalian L1 elements, for which mobility competent loci were identified (27–34). These donor L1 loci were shown to carry unmethylated promoters, lineage-specific transcription factor motifs or deletions of binding sites for repressive factors, all contributing to their activity (22–24,35,36).

In contrast to L1 elements, identification of somatically active loci of LTR retrotransposons and an understanding of their regulation have been lagging behind. This is in part because of the fact that no replication-competent LTR elements have been identified in the human genome. However, their transcription in different tissues is substantial (16–18) and they continue to mobilize in other species, including model organisms such as mouse (37) or *Drosophila* (38,39). It is generally believed, that expression of LTR retrotransposons is determined by the 5’UTR region, including the LTR sequence itself (40–44). Indeed, sequence variations of these regulatory regions were proposed to explain transcriptional activity and tissue or cell-type specific patterns of expression of LTR retroelements (16–19). However, other levels of regulation are likely to play a role and they may be uncovered through identification and careful analysis of somatically active LTR-element loci.

In our previous work we showed that retroelements are expressed and mobile in *D. melanogaster* intestinal tissue and that somatic transposition can lead to gene inactivation via LTR-element insertion (39). Here, using long-read DNA sequencing, we further characterize this mobility in healthy somatic tissues isolated from flies of different ages. Importantly, we uncover a donor locus of a *rover* sub-family of endogenous retroviruses (ERV). ERVs constitute a sub-class of LTR retrotransposons, which acquired an *envelope* (*env*) open reading frame (ORF), in addition to the *gag* (capsid) and *pol* ORFs carried by all LTR retroelements (45,46). We demonstrate that the activity of the somatically active *rover* retroviral locus is driven by its genomic *cis* regulatory elements, providing new insight into somatic regulation of LTR/ERV elements by their genomic environment.

## Results

### The genomic landscape of fixed, rare and somatic TE insertions

To gain new insight into somatic mobile element activity, we explored our previously-published long-read DNA sequencing datasets (39) and generated new, complementary libraries (Supp. Table 1). Using Oxford Nanopore Technology (ONT), we sequenced DNA isolated from pools of *Drosophila* heads or intestines, using one of our previously characterized genetic backgrounds with documented TE mobility in the gut (*ProsGAL4>UAS-GFP* (39,47), hereafter abbreviated as *ProsGFP*). Of note, the *ProsGFP* flies are wild-type for the known TE controlling pathways (39). For each tissue type, we sequenced in two replicates DNA libraries from young (5-7-day-old), mid-aged (25-30-day-old) and old (55-60-day-old) female flies, obtaining on average a sequencing depth of 50x (Supp. Table 1).

First, we aimed to precisely characterize the TE landscape of the studied genotype, focusing on full-length copies (that comprise more than 70% of the respective consensus sequence, see Materials and Methods section). To do this, we analyzed long-read DNA sequencing libraries for TE insertions annotated in the dm6 reference genome and non-reference TE insertions present in the genome of the *ProsGFP* strain. Among 4444 full-length TEs (Supp. Table 2), 1685 (38%) were also found in the dm6 reference genome (***Figure 1A***). The remaining 62% were non-reference. We genotyped all detected full-length TE insertions based on their: 1) presence in the reference genome; 2) ONT read ratio (number of supporting *vs* opposing reads), 3) tissues in which their insertion was detected, and 4) germline activity of their respective sub- families (details in Materials and Methods). We further used our 31 previously published Illumina DNA sequencing libraries of head and intestine tissues coming from individual flies of the *ProsGFP* genetic background (39) to validate genotype assignment. As expected, reference TEs had in general ONT read ratios close to 1 (***Figure 1B***), reflecting their homozygous status. We subdivided non-referenced TEs into 4 categories: “Fixed non-reference”, “Rare”, “Singleton”, and “Ungenotyped”, described further below.

**Figure 1.**
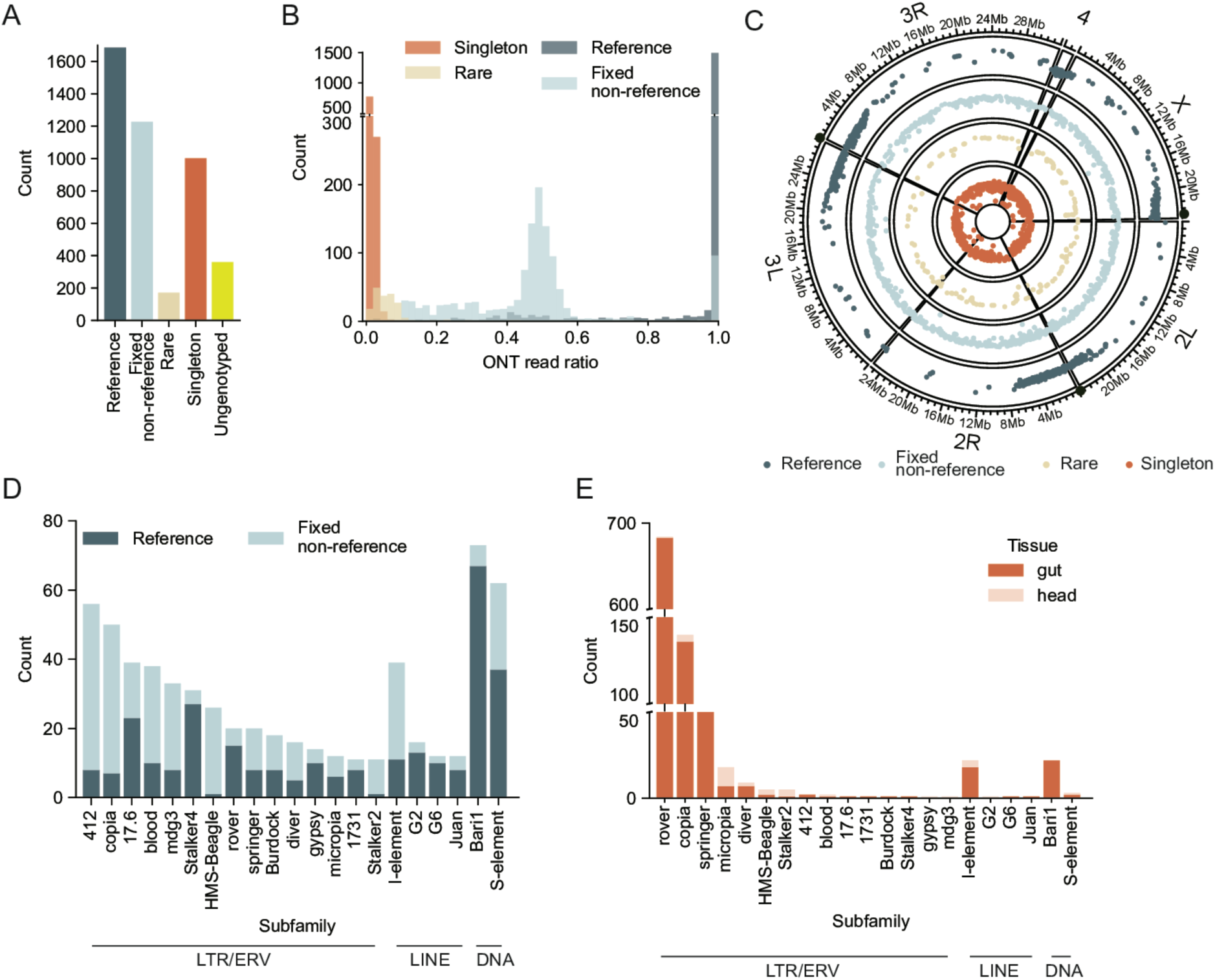
The landscape of full-length fixed and somatic TE insertions using long-read DNA sequencing. (**A**) Numbers of full-length TE insertions in the *ProsGFP* genome categorized in the “reference” (dark blue), “fixed non-reference” (light blue), “rare” (light brown) and “singleton” (somatic, orange) genotypes, as well as the “ungenotyped” insertions. (**B**) Distribution of ONT read ratios (number of supporting *vs* opposing reads) for TE insertions categorized as “reference” (dark blue), “fixed non-reference” (light blue), “singleton” (somatic, orange) and “rare” (light brown). (**C**) Genome-wide distribution of the detected “reference” (dark blue), “fixed non-reference” (light blue), “singleton” (orange) and “rare” (light brown) TE insertions on the *Drosophila* chromosome arms. Black circles indicate positions of centromeres. (**D**) Numbers of full-length “reference” (dark blue) and “fixed non-reference” (light blue) insertions of the different TE sub-families. Only the sub-families which contributed somatic insertions are plotted. For all other TE sub-families, see Supplementary Figure 1C. (**E**) Numbers of somatic insertions (“singletons”) of different TE sub-families recovered from the gut (dark orange) or head (light orange) DNA libraries.

We first characterized insertions that were detected in at least one ONT sample of each tissue with ONT read ratio higher than 0.1 (1227 insertions in total), which were assigned the “fixed non-reference” genotype (***Figure 1A-B***). ONT read ratios of these insertions peaked at 0.5 or 1, likely reflecting predominantly germline heterozygous and homozygous TE insertions, respectively (***Figure 1B***, ***Supplementary Figure 1A***). A high number of non-reference insertions was consistent with previous reports on the high variation in germline TE composition in *D. melanogaster* laboratory strains (38,48). Only 60% (747 out of 1227) of “fixed non-reference” insertions were detected in our previously published Illumina DNA sequencing libraries from the same genetic background (39), 98% of which were classified as germline TE insertions. Moreover, the distribution of Illumina variant allele frequencies (VAFs) and population frequencies of these insertions supported the idea that most of the “fixed non-reference” insertions are heterozygous insertions fixed in the strain (***Supplementary Figure 1A***). 171 insertions with ONT read ratio lower than 0.1 that were detected in at least one sample of each tissue were assigned “Rare” genotype (***Figure 1A and B***). Low ONT read ratios suggested that they represented rare germline insertions, present only in some individuals (as in (49)). Only 25 of them (15%) were also detected in the short-read samples. Together, this analysis highlights the high variation in TE composition between the reference genome and the investigated genetic background and the advantage of long-read sequencing to comprehensively detect TE sequences.

We then focused on somatic insertions. We defined these as in our previous study (39), as insertions being supported by a single read present in only one sequencing library (“singleton”) that had a target site duplication (TSD) as a footprint of a true transposition event. In addition, we excluded sub-families active in the germline (***Supplementary Figure 1B***). We detected 1001 of such “singleton” insertions (***Figure 1A and B***). We have previously provided strong evidence that “singletons” can confidently be considered as true somatic insertions in this experimental and computational setup (39) and we further introduced additional filtering steps to minimize potential false positives (see Material and Methods for details).

Finally, the insertions that did not meet the criteria for “fixed non-reference”, “rare” and “singleton” genotypes were categorized as “ungenotyped” insertions. We detected 360 such insertions (Figure 1A). These are insertions either with ambiguous set of supporting samples, or singletons lacking TSD footprint, or singletons from the sub-families active in the germline. They could represent artifacts, very rare germline insertions, somatic insertions including embryonic and developmental events, or a mixture of these three, which we cannot confidently distinguish.

Having described the TE landscape of the *ProsGFP* strain, we then aimed to investigate the genomic TE distribution (***Figure 1C***). In agreement with previous reports (38,50), fixed referenced insertions were found enriched in pericentromeric, gene-poor chromosome regions, and on the mostly heterochromatic chromosome 4. Conversely, fixed non-referenced insertions were present throughout all chromosome arms, without any significant “hot-spots” similarly to the rare insertions. Accumulation of fixed reference insertion in pericentromeric regions is likely an effect of a negative selection acting on these, likely evolutionarily “older”, germline insertions. In contrast, fixed non-reference and rare TEs possibly represent more recent germline insertions. Finally, consistent with our previous data (39), singleton (somatic) TE insertions were distributed genome-wide, similarly to the germline non-refence and rare insertions.

Last but not least, we examined TE sub-families represented in each genotype. Full-length fixed reference and non-reference insertions were recovered for 120 out of 126 known *Drosophila* TE sub-families, including Class I retrotransposons (the most abundant in terms of copy number and total sequence in the reference genome (38,51)) and Class II DNA elements (***Figure 1D*** and ***Supplementary Figure 1C***). Rare insertions were represented by 34 sub-families (***Supplementary Figure 1D***).

Consistent with what we reported previously and in contrast to the fixed insertions, only a few TE sub- families were represented among singleton insertions (***Figure 1E***). A great majority of singletons (904 out of 1001) were retrotransposon insertions of three sub-families: ERV element *rover* (684 singletons), LTR element *copia* (140 singletons) and ERV element *springer* (80 singletons.) We also recovered 22 singleton insertions of a non-LTR, LINE-like retrotransposons *I-element*, as well as 22 insertions of *Bari1*, a sub- family of DNA transposons. Notably, 965 (96%) of detected singletons were found in the libraries obtained from the intestines, while only 36 (4%) were head-specific insertions (***Figure 1E***).

Taken together, the analysis of long-read DNA sequencing datasets enabled in-depth characterization of the full-length fixed TE insertions as well as detection of somatic events specific for one tissue type and limited to a small number of somatically active TE sub-classes. Since significant somatic mobility was detected only in the DNA sequencing libraries from the gut samples, we further focused on this tissue.

### Aging is not associated with important increases in somatic *de novo* TE insertions

In different species, heterochromatin relaxation and transcriptional de-repression of DNA repeats, including TEs, are associated with aging (52–54). Therefore, we next asked if significant changes in somatic TE activity could be detected between young, mid-aged or old tissues. To this end, we first carried out expression analysis using short-read RNA-seq data from young (39) as well as aged (this study) midguts to establish TE expression levels (***Figure 2A***). In agreement with previously published reports from the fly intestine (55,56), we detected increased transcript levels of selected TE sub-families in old guts, including, retrotransposons *copia*, and *springer*. Nevertheless, age-related upregulation in transcript levels was not widespread throughout TE sub-families (14 upregulated TE sub-families) and a comparable number of TEs was also significantly down-regulated (8 TE sub-families). Out of the three LTR/ERV sub- families identified as the most mobile (***Figure 1G***), *copia* and *springer* showed increased transcript levels, while *rover* transcripts were not significantly changed. Moreover, neither the expression levels nor the extent of the age-related deregulation correlated with the overall TE mobility (***Supplementary Figure 2A-C***).

**Figure 2.**
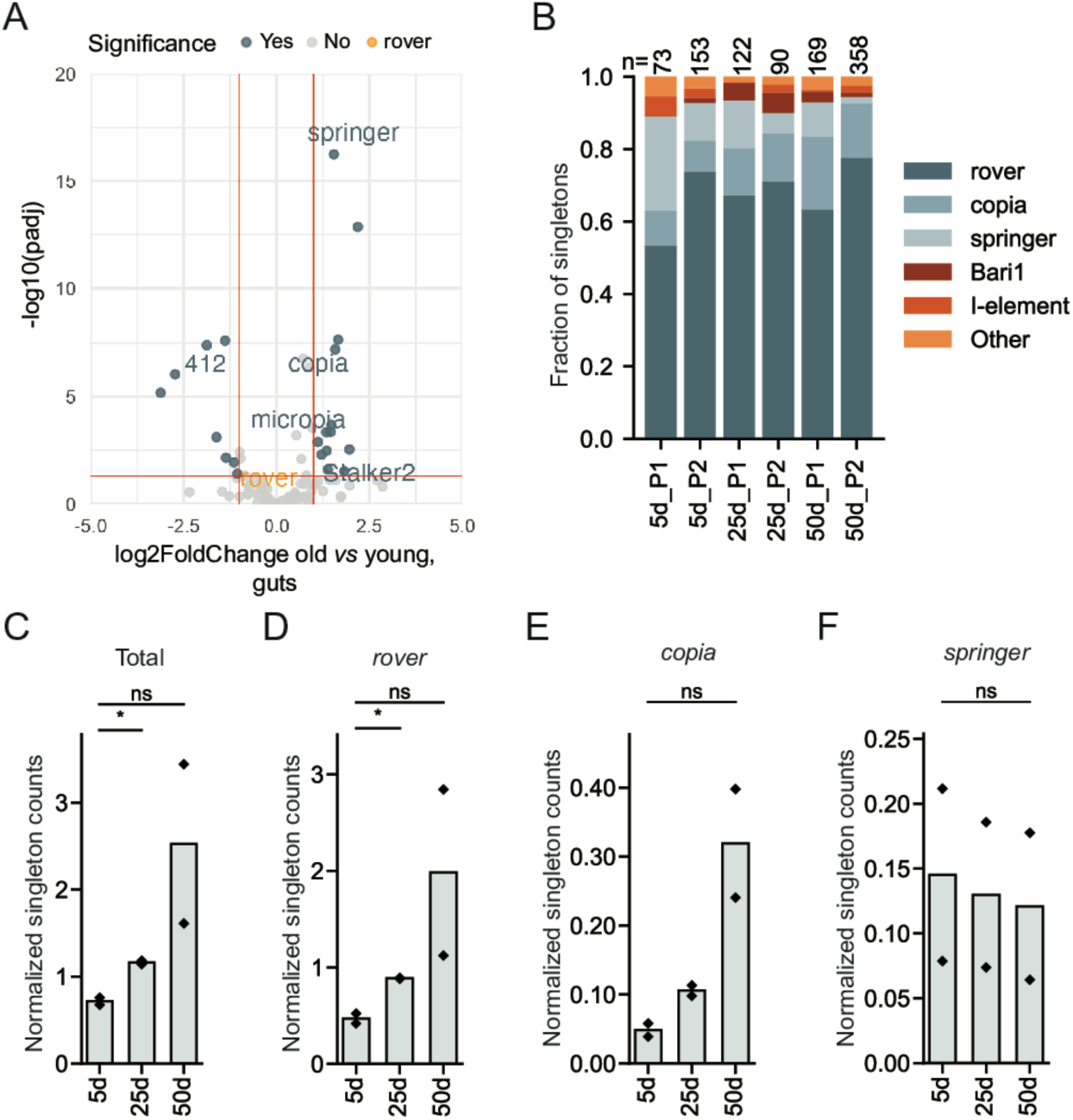
Somatic TE expression and mobility during aging. (**A**) Volcano plot illustrating changes in TE transcript levels upon aging in guts of the *ProsGFP* genotype. Data is from short-read RNA-seq. Significantly up- or down-regulated TEs are indicated in blue and TE sub-families which were detected as mobile are labeled. (**B**) Sub-family distribution of somatic TE insertions detected in the gut DNA libraries from young (5 days), mid-aged (25 days) and old (50days) flies. P1 and P2 are two biological replicates (pool 1 and 2). n= total number of “singletons” in each pool. (**C-F**) Normalized counts of somatic insertions detected in libraries obtained from gut tissues of flies with different ages (5, 25 or 50 days). Plotted are all TE sub-families (**C**), as well TE sub-families contributing the most *de novo* insertions: *rover-ERV* (**D**), *copia-LTR* (**E**) and *springer-ERV* (**F**). ns, not significant; * adjusted p-value < 0.05 (one-way ANOVA test).

Since we have previously shown that TE transcript levels reflect poorly the actual TE mobility (39), we next focused on comparing somatic TE *de novo* insertions between the age groups. The contribution of TE sub- families to the total number of singleton insertions in each library was not considerably different between libraries from young, mid-aged and old guts, with *rover*, *copia* and *springer* LTR/ERV-type elements dominating in all samples (***Figure 2B***). ONT sequencing libraries can significantly vary in terms of yield and read lengths, making direct comparison of the raw singleton counts between libraries difficult. Thus, we normalized the raw singleton counts for sequencing depth considering read length distributions in each library. By doing so, we observed a moderate increase with age in the total, *rover-ERV, and Bari1-DNA* normalized singleton counts (***Figure 2C and E***, and ***Supplementary Figure 2E***). The significant change in the total and the *rover-ERV* counts could be detected between samples from young and mid-aged flies, but not between young and old individuals, likely stemming from the high variability between the two replicates in the latter. No significant changes between age groups were detected when singletons of *copia*, *springer or I-element* sub-families were counted (***Figure 2D and F***, and ***Supplementary Figure 2D***). Thus, only minor changes in somatic TE insertion loads could be detected between DNA samples form young and aged gut.

We then sought to determine if similar results were obtained for the fly head samples. In the head transcriptomes, expression of 18 TE sub-families was upregulated in aged tissues, and three sub-families were downregulated (***Supplementary Figure 2F***). Among the significantly deregulated TEs, *copia* (upregulated) and *mdg3* (downregulated) were mobile in the head tissue according to our analysis (***Supplementary Figure 2G***). However, the association of the transcriptional deregulation and the mobility could not be reliably tested owing to the low numbers of the detected singletons. Thus, expression of few TE sub-families was changes in aged fly heads of the investigated genetic background, but the detected *de novo* TE insertions were marginal in all age groups.

In conclusion, even though transcript levels change for selected TE sub-families in the aging fly gut and head tissues, we could not observe important age-related triggering of TE mobility both in terms of insertion numbers or active TE sub-families.

### Mobility and expression of the *rover*-LTR/ERV sub-family is restricted to one donor locus

To gain insights into the mechanisms of retrotransposon activation in the soma, we then sought to identify potential donor retrotransposon loci responsible for the observed somatic mobility. We focused our analysis on the *rover-ERV* sub-family that accounted for almost 70% of all somatic insertions. Taking advantage of ONT reads spanning full insertion length between genomic breakpoints, we extracted sequences of *rover* singletons, representing somatic insertions, and aligned them to the *rover* consensus sequence (***Figure 3A***). The genome of the *ProsGFP* strain contains 15 nearly full-length reference and 5 fixed non-reference *rover* copies. We created consensus sequences for the fixed *rover* insertions using ONT reads and aligned them to the published *rover* consensus sequence (https://github.com/bergmanlab/drosophila-transposons). Visual inspection of the alignments of the fixed and singleton insertions showed that 1 fixed, non-reference and 11 reference insertions had large structural variants that were not present in the singleton insertions (***Supplementary Figure 3***), leaving 8 full-length potential candidates to be the donor locus (***Figure 3A and B***). Only 2 out of the 8 candidate loci contained the small structural variants present in the singletons. Moreover, almost all singletons showed support for the single nucleotide polymorphisms (SNPs) present in only one of these two *rover* loci (96% singletons on average per SNP, Supplementary Table 3). The discrepancy from 100% was within ONT sequencing error rate and none of the singletons had the full set of SNPs that supported the second locus, suggesting that the donor locus is on 2^nd^ chromosome at position chr2R:14487730-14487747 (***Figure 3B***). This locus is in the anti-sense orientation within the first intron of a *PRAS40* gene (***Figure 3C***) and is not present in the dm6 reference genome. Thus, we concluded that the observed somatic mobility of the *rover- LTR/ERV* sub-family in the intestinal tissue is limited to one donor locus, here named *rover-2R:14M*.

**Figure 3.**
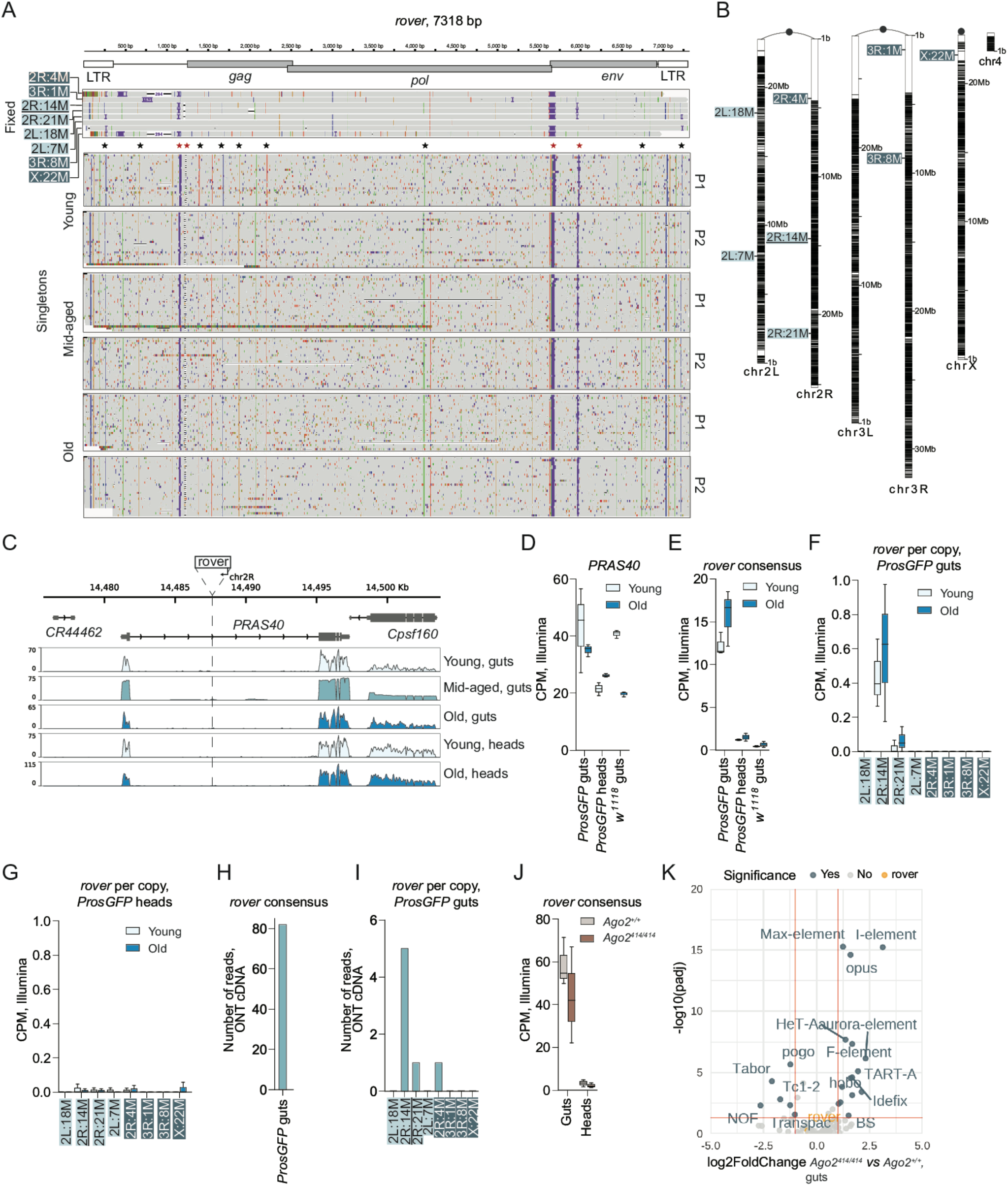
Somatic activity of the *rover-LTR/ERV* family is limited to one donor locus. (**A**) Alignment of the fixed *rover* copies present in the analyzed genome and a selection of somatic (singleton) *rover* insertions detected in different long-read DNA-seq libraries to the published *rover* consensus sequence. The fixed copies (covered by many reads) are named following their position in the genome and each line represents a consensus sequence of the copy. For somatic insertions, each line represents a single read covering an insertion. Fixed reference copies are marked with dark blue and white lettering, and fixed non-reference copies with light blue. The identified donor *rover-2R:14M* copy is underlined. Regions with structural variants and single nucleotide polymorphisms that distinguish the donor *rover-2R:14M* copy from other *rover* copies are highlighted with red and black stars respectively. (**B**) Schematic representation of the *Drosophila* chromosomes with the positions of fixed *rover* copies in the *ProsGFP* genome. Fixed reference copies are marked with dark blue and white lettering, and fixed non-reference copies with light blue. The identified donor *rover-2R:14M* copy is underlined. (**C**) Position of the donor *rover-2R:14M* locus on the chromosome 2R within the intron of the *PRAS40* gene (upper panel). RNA-seq coverage of the *PRAS40* locus showing expression of the gene in libraries obtained from gut or head tissues from flies of different ages (lower panels). Young and old gut, and young and old head data are from short-read RNA-sequencing. Mid-aged gut data are from ONT long-read cDNA sequencing. (**D-E**) Normalized expression levels (CPM, counts per million) in gut and head transcriptomes of the *ProsGFP* genotype (with the *rover-2R:14M* insertion) and in gut transcriptomes of the *w^1118^* control genotype (without the *rover-2R:14M* insertion). Quantified are transcript levels of *PRAS40* (**D**) and of *rover* upon read mapping to the consensus sequence (**E**). Data is from short-read RNA-seq. (**F-G**) Normalized expression levels (CPM, counts per million) of *rover* upon copy-specific read mapping in transcriptomes from guts (**F**) and heads (**G**) of the *ProsGFP* strain. Data is from short-read RNA-seq. Fix reference copies are marked with dark blue and white lettering, and fixed non-reference copies with light blue. The identified donor *rover-2R:14M* copy is underlined. (**H-I**) Number of long reads mapping to the *rover* consensus (**H**) and to the different *rover* copies present in the genome (**I**) in long-read transcriptomes from the *ProsGFP* background. Data is from ONT cDNA sequencing. Fix reference copies are marked with dark blue and white lettering, and fixed non-reference copies with light blue. The identified donor *rover-2R:14M* copy is underlined. (**J**) Normalized *rover* expression levels (CPM, counts per million) in gut and head transcriptomes of the *Ago2* mutant (*rover-2R:14M^+/+^; Ago2^414/414^*) and the *rover-2R:14M^+/+^; Ago2^+/+^* control flies. Reads were mapped to the *rover* consensus sequence. (**K**) Volcano plot illustrating changes in TE transcript levels in *Ago2* mutant (*rover-2R:14M+/+; Ago2^414/414^*) guts as compared to *rover- 2R:14M^+/+^;Ago2^+/+^* control tissues. Both genotypes carry the *rover-2R:14M* active copy. Significantly up- or down-regulated TE sub-families are indicated in blue and labeled.

As the first prerequisite for retrotransposon mobility is its transcription, we next aimed to confirm the expression of the *rover-2R:14M* and check if other, non-mobile, *rover* copies were also expressed in the tissue. Aligning the short transcriptomic reads to the *rover* consensus sequence confirmed *rover* expression in the gut tissue, regardless of the age of the flies, while minimal expression was detected in the head samples (***Figure 3C and E***). Additionally, we did not detect *rover* transcripts in the gut samples of the *w^1118^* genetic background (commonly used wild-type stock), which did not carry the *rover-2R:14M* copy (***Figure 3E***). We then attempted to quantify per-copy expression (***Figure 3F and G***). To address sequence similarity among rover copies, we extracted the reads that aligned to the *rover* consensus sequence and re-aligned *them* with stringent parameters to the consensus sequences of all fixed *rover* copies in the *ProsGFP* genome, discarding multimapping reads. This approach, which did not provide absolute per-copy expression levels but instead showed expression levels relative to other copies, revealed that the *rover-2R:14M* locus was the only full-length locus of the *rover* sub-family expressed in the gut, consistent with our evidence above from rover mobility. Finally, in order to further validate this result, we additionally performed long-read ONT cDNA sequencing on midguts isolated from mid-aged flies, because short-read sequencing has important limitations in the RNA-seq analysis of repeat sequences (21) (***Figure 3H and I***). Consistent with the short-read datasets, we saw evidence for a predominant expression of the donor *rover-2R:14M* locus (***Figure 3I***), suggesting that other *rover* loci were not, or very lowly expressed in the gut tissue.

We then asked how *rover-2R:14M* expression compared to the expression of the *PRAS40* gene in which *rover-2R:14M* is embedded. Importantly, *rover-2R:14M* expression did not fully correlate with that of *PRAS40*: *PRAS40* was found to be expressed in both gut and head tissues (***Figure 3D***) in contrast to *rover-2R:14M* expressed in the gut, suggesting distinct transcriptional regulation. In addition, the presence or absence of *rover-2R:14M* within *PRAS40*, did not alter *PRAS40* expression patterns in either the head or the gut, as seen by comparing the *ProsGFP* background (*rover-2R:14M* positive) to the *w^1118^* background (*rover-2R:14M* negative) (***Figure 3C and D***). Moreover, the *rover-2R:14M* insertion did not cause mis-splicing of the *PRAS40* gene as shown by the ONT cDNA sequencing data (***Supplementary Figure 4A***). However, we observed a single transcript in the ONT cDNA sequencing data, 5’ end of which was located in the *rover-2R:14M* 3’LTR and which contained the downstream *PRAS40* intronic sequence. This indicated that *rover-2R:14M* locus could potentially enable weak aberrant transcription in the gut tissue (***Supplementary Figure 4A’***). Together this analysis hinted that the *rover-2R:14M* donor locus is expressed in the gut without affecting the transcription of the gene it is embedded in.

The control of TE transcript levels in the fly non-gonadal tissues is mostly achieved by the endogenous short interfering (siRNA) pathway, responsible for sequence-specific transcript degradation (57–59). We have previously demonstrated that *rover* mobility in the *ProsGFP* flies occurs despite the presence of siRNAs against *rover* in the gut (39), suggesting that post-transcriptional silencing of *rover* may not be efficient in this genetic background. To further address if the identified *rover-2R:14M* locus is regulated by the siRNA pathway, we then performed transcriptome analysis in tissues isolated from flies carrying the *rover-2R:14M* locus and mutant for *Argonaute 2* (*Ago2*), deficient in the siRNA pathway, as well as their isogenic controls (***Figure 3J and K***). In both genetic backgrounds *rover* transcripts were consistently detected only in the gut RNA-transcriptomes and not in the heads. Importantly, *rover* transcript levels were not changed in the Ago2 homozygous mutant tissues (***Figure 3J***), even though we detected differential expression of other TE sub-families in the gut and the head transcriptomes (***Figure 3K and Supplementary Figure 5***), as previously reported in *Ago2* mutant tissues (57–59). Altogether, these data suggest that the identified donor *rover-2R:14M* locus expressed in the fly gut is not post-transcriptionally regulated by the siRNA pathway.

Interestingly, in a concomitant independent study using a different fly genetic background for the detection of *de novo* TE insertions, activity of the *rover* sub-family was also reported and classified as likely somatic (60). Using their data, we have performed the same type of sequence comparison of the fixed *rover* copies and the *de novo* singleton insertions from that study, and found that the *rover-2R:14M* insertion was also present in the investigated genetic background and likely acted as a source locus for the great majority of *de novo* somatic insertions (***Supplementary Figure 6***).

Thus, we identified a non-reference polymorphic “hot” *rover-2R:14M* LTR/ERV retroelement locus, transcribed in the fly intestinal tissue and serving as a donor locus for *de novo* somatic insertions.

### The donor *rover-2R:14M LTR/ERV* locus is located in permissive chromatin

We then aimed to understand which mechanisms could underlie the somatic activity of the *rover-2R:14M* locus. Since LTR/ERV retrotransposons are believed to carry their own regulatory sequences within the LTR sequence itself and the larger 5’UTR region proceeding the LTR (40–44), we first asked if the transcriptional activity of the *rover-2R:14M* locus could be explained by sequence variation private to the locus within the transposable element sequence. We analyzed the 5’ LTR sequences, sites of transcriptional initiation of LTR-type elements, of the donor locus and two non-active *rover* loci (*2R:21M* and *2L:18*), which had the highest sequence similarity to the donor locus. We first assessed the presence of the sequence motifs for the Polymerase II (PolII) transcription initiation complex, namely TATA-box, initiator element (Inr), downstream promoter element (DPE) and motif ten element (MTE), as LTR elements are believed to be PolII transcribed (61) (***Figure 4A***). We did not notice significant differences between the full-length *rover* copies that could explain gut-specific expression. We then extended the analysis to an *in silico* analysis of putative transcription factor (TF) binding sites within the first 2 kb of the internal *rover* sequences (***Figure 4B*** and ***Supplementary Figure 7***). Again, we were unable to find motifs that were different between the active *rover-2R:14M* locus and non-expressed loci, implying that internal sequence variation could not account for the differences in the activity of the compared *rover* loci.

**Figure 4.**
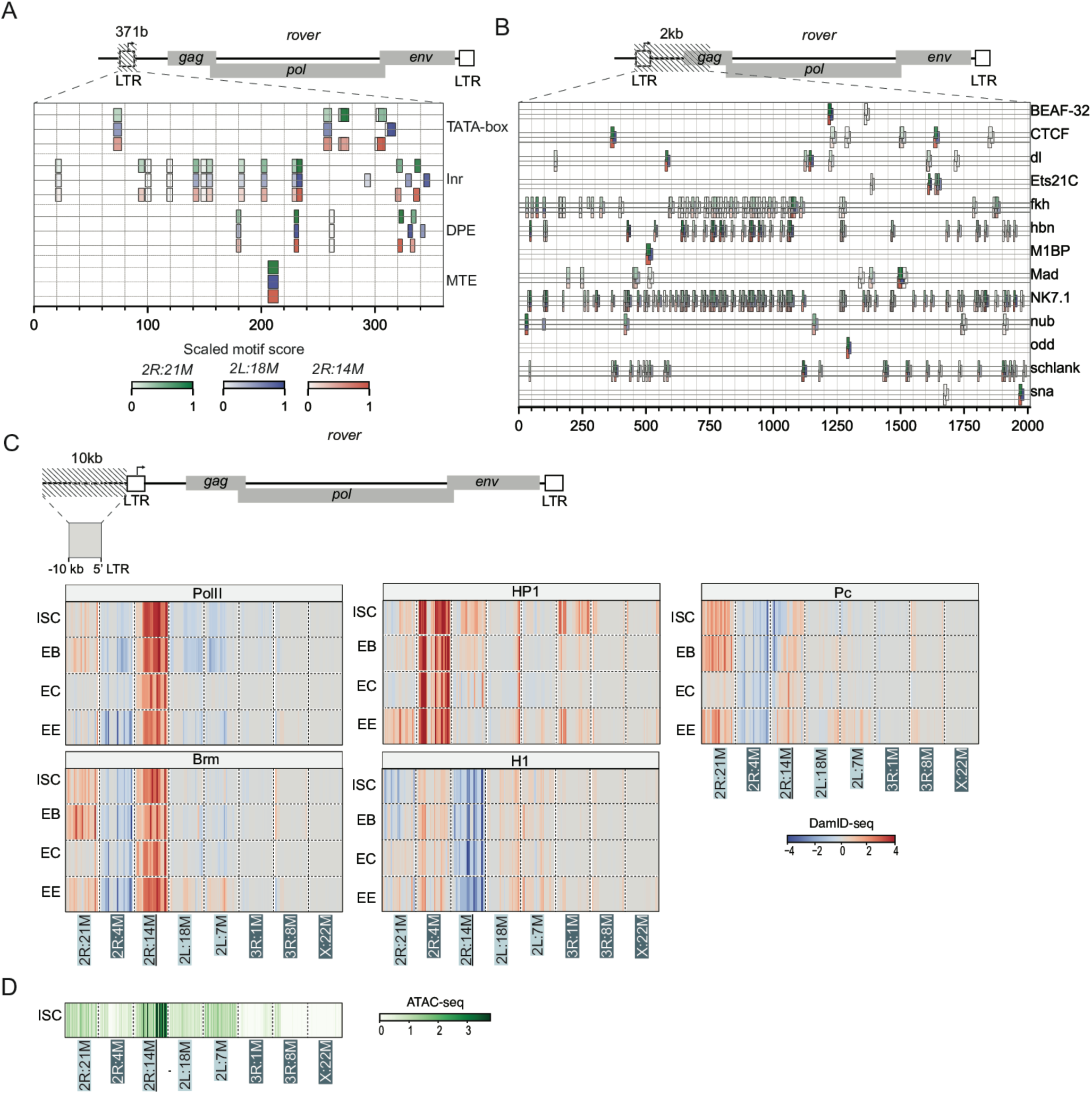
The donor *rover-2R:14M* locus is located in permissive chromatin. (**A**) *In silico* detection of DNA motifs for Polymerase II (PolII) transcription initiation complex (TATA-box, Inr, DPE and MTE) in the 5’LTR region of the donor *rover-2R:14M* copy (red) and two non-active copies with the highest sequence similarity (*2R:21M*, green and *2L:18*, blue). (**B**) *In silico* search for transcription factor (TF) binding sites within the first 2 kb of the internal sequences of the donor *rover-2R:14M* copy (red) and two non-active copies with the highest sequence similarity (*2R:21M*, green and *2L:18*, blue). Only TFs highly expressed (RPKM>10) in the gut lineage are plotted. (For all TFs, see Supplementary Figure 7). (**C**) Gut cell type-specific heatmaps of DamID chromatin profiles upstream (10 kb) of the active *rover-2R:14M* copy and 7 inactive full-length *rover* loci present in the investigated genome. 5 chromatin binding factors are shown: Polymerase II (PolII), Brahma (Brm), Polycomb (Pc), heterochromatin protein 1a (HP1) and histone H1 (H1). Intestinal Stem Cells (ISC) and Enteroblasts (EB) are gut progenitor cells. Enterocytes (EC) and Enteroendocrine cells (EE) are differentiated cell types. Fixed *rover* reference copies are marked with dark blue and white lettering, and fixed non-reference copies with light blue. The identified donor *rover-2R:14M* copy is underlined. (**D**) Heatmaps of intestinal stem cell (ISC) ATAC-seq profiles upstream (10 kb) of the active *rover-2R:14M* copy and 7 inactive full-length *rover* loci present in the investigated genome. Fixed reference copies are marked with dark blue and white lettering, and fixed non-reference copies with light blue. The identified donor *rover-2R:14M* copy is underlined.

We thus reasoned that *rover-2R:14M* transcriptional activity (and mobility) might be conferred by its genomic position rather than its internal sequence. Given that, we inspected chromatin environment upstream (10 kb) of the active and 7 inactive full-length *rover* loci present in the investigated genome, making use of the published gut cell type-specific DamID (DNA adenine methyltransferase identification) datasets profiling 5 chromatin binding factors associated with “active” or “silent” chromatin states (62,63) (***Figure 4C***). Only two *rover* loci (*2R:21Mb* and *2R:14Mb*) were located in “active” regions bound by Polymerase II (PolII) and Brahma (Brm), with the upstream region of the donor *2R:14Mb* locus showing the strongest binding in all intestinal cell types. The upstream regions of the remaining loci were either not bound at all or bound by the heterochromatin factors heterochromatin protein 1a (HP1), histone H1 (H1) or Polycomb (Pc). Similarly, when the same regions were inspected using published ATAC-seq (Assay for Transposase-Accessible Chromatin) data (63), the upstream region of the donor *rover-2R:14M* locus showed the highest accessibility signal (***Figure 4D***), and thus the most “open” chromatin environment.

Altogether, this analysis indicated that the epigenomic environment, rather than copy-specific sequence variants, is likely responsible for the *rover-2R:14M* activity in the gut tissue.

### *rover-2R:14M* expression is driven by its upstream genomic sequence

To experimentally test the transcriptional regulation of the active *rover* copy, we engineered expression reporters where the *rover-2R:14M* 5’UTR and upstream genomic sequences were placed in front of a *lacZ.NLS* (NLS, nuclear localization signal) reporter gene (***Figure 5A***). We inserted the reporters into the fly genome using two different landing sites on the X and the 3rd chromosomes. The sites were chosen based on their similarity in the DamID chromatin landscapes to the original *rover-2R:14M* genomic location (“active” chromatin in all gut cell types, ***Figure 4C*** and ***Supplementary Figure 8***).

**Figure 5.**
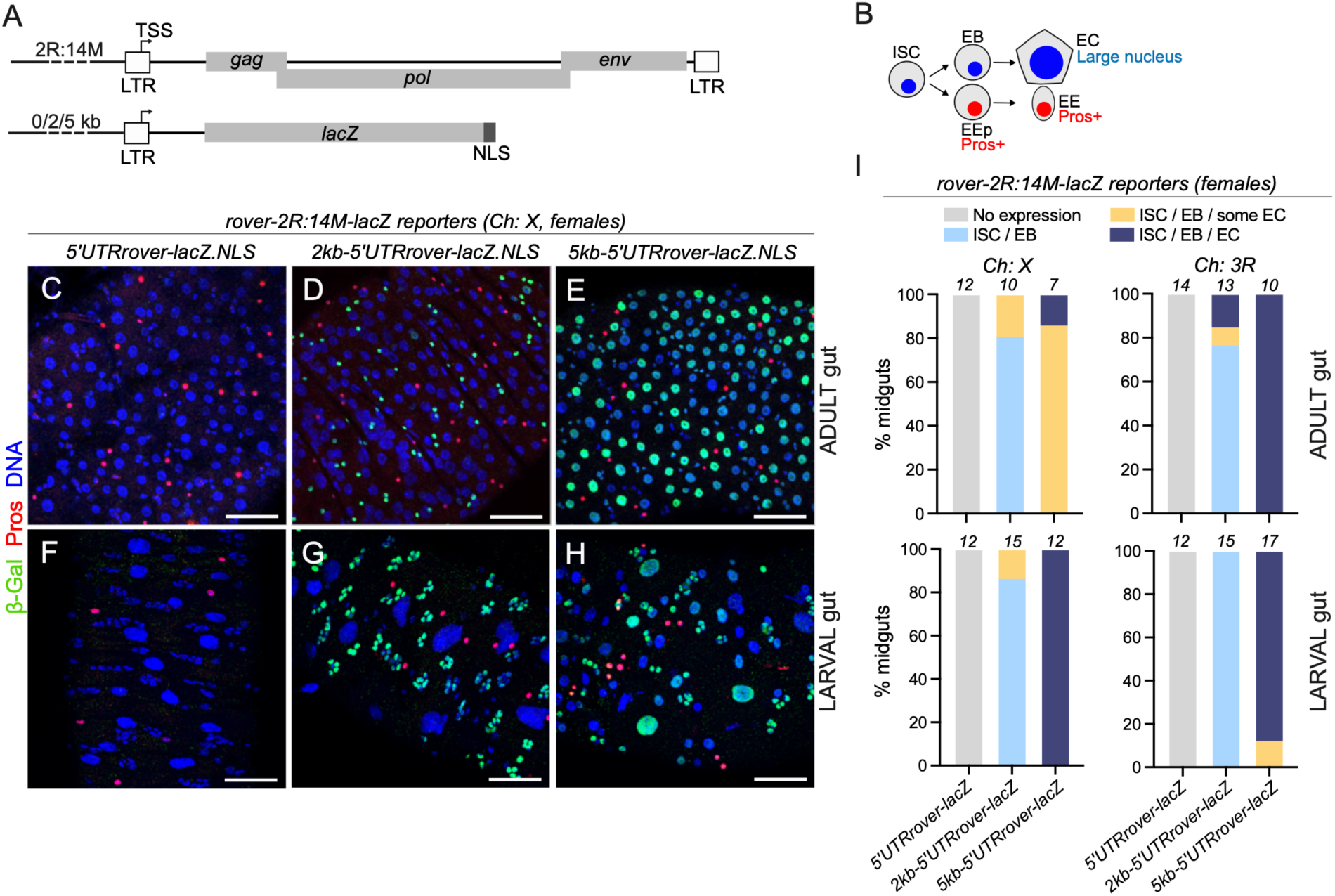
Upstream genomic sequence drives *rover-2R:14M* expression in the gut. (**A**) Schematic representation of the *in vivo rover-2R:14M* reporter constructs. lacZ reporter was placed downstream of the *rover-2R:14M* 5’UTR region replacing all *rover* native open reading frames. 0, 2 or 5 kb of the upstream genomic sequence was also added. (**B**) Schematic representation of the *Drosophila* intestinal lineage. Intestinal stem cells (ISCs) give rise to committed progenitors: enteroblasts (EBs) and enteroendocrine precursors (EEp). The progenitors then differentiate into enterocytes (ECs) and enteroendocrine cells (EEs). The EEp and EE express transcription factor Prospero (Pros). The ECs are distinguished by a large, polyploid nucleus. In the larval gut, the stem cells, named adult midgut precursors (AMPs), form distinct clusters. (**C-H**) Representative images of female midguts carrying *rover-2R:14M-lacZ* reporter constructs inserted on the X chromosome stained for β-galactosidase (β-gal, green), Prospero (Pros, red) and DNA (blue). Adult (C-E) and larval (**F-G**) tissues are shown. Scale bar: 50 μm (**I**) Quantifications of the expression patterns of *rover-2R:14M-lacZ* reporter constructs in female adult and larval midguts. Numbers of midguts scored are indicated above each bar.

To begin to define the patterns or the reporter activity *in vivo*, we first tested *lacZ* expression in the female gonads, where similar reporters for other retrotransposon families were already used (64–66). As previously shown, in ovaries with functional small RNA-driven TE silencing, we detected no or very weak expression of *lacZ* under the control of the 5’UTR of *rover-2R:14M* (***Supplementary Figure 9A-B’*** and ***E- F’***). However, *lacZ* expression was readily detected upon knock-down of the piRNA pathway with a germline- (***Supplementary Figure 9C-D’***) or somatic follicle cell-specific drivers (***Supplementary Figure 9G-H’***). This confirmed that, as expected, in female reproductive tissues, the *rover-2R:14M* reporter can be efficiently transcribed, but is silenced by the piRNA pathway.

To gain insight into *rover-2R:14M* expression in the intestine, we then examined the lacZ reporter expression in this tissue in the adult stage, as well as during development, in the 3^rd^ instar larval stage. The adult and the larval gut consist of progenitors and differentiated cell types, which can be distinguished by the expression of cell-type specific markers and by the nuclear size (***Figure 5B***). *LacZ* was not expressed in the gut when driven only by the 5’UTR region of *rover-2R:14M* (***Figure 5C, F*** and ***I***). However, by extending the regulatory region with 2 kb or 5 kb of the upstream genomic sequence, we could achieve reporter expression in adult (***Figure 5D-E*** and ***I***) and larval intestinal tissues (***Figure 5G-H*** and ***I***), even without interfering with the TE silencing pathways. Patterns of reporter expression were different depending on the length of the upstream regulatory sequence. 2 kb of the upstream region drove *rover- 2R:14M* reporter expression mostly in gut progenitor cells: stem cells (ISCs) and enteroblasts (EBs) (***Figure 5D, G*** and ***I***, unmarked small diploid cells). Increasing the length of the sequence to 5 kb allowed for broader reporter expression patterns in progenitors as well as differentiated absorptive enterocytes (ECs, large polyploid nuclei) (***Figure 5E, H*** and ***I***). We did not detect reporter activity in the secretory enteroendocrine (EE) cells, marked by the expression of the Prospero (Pros) transcription factor. These expression patterns were largely consistent between females and males (***Supplementary Figure 10***). Notably, we obtained similar results with both insertion sites tested (***Figure 5I***), suggesting that, at least for the two insertion sites tested by us, reporter expression was not influenced by the genomic position.

Taken together, these results support the idea that *rover-2R:14M* expression in the fly gut may be driven in *cis* by its upstream genomic sequence.

### Escargot transcription factor regulates *rover-2R;14M* expression

To better understand how the upstream genomic region could allow for *rover-2R:14M* expression in the gut, we next analyzed published chromatin immunoprecipitation (ChiP-seq) data (ModEncode Project, (67)) to identify transcription factor binding sites present in the region (***Figure 6A***). Indeed, many transcription factors were found to bind to the region, particularly within the first 2 kb upstream of the *rover- 2R:14M* insertion. We sorted the transcription factors based on their expression in the adult gut intestinal stem cells (***Figure 6A***), enteroblasts or differentiated enterocytes (***Supplementary Figure 11***) using previously published transcriptomic data (68). Based on this analysis we identified a set of candidate factors, that could regulate *rover-2R:14M* expression through the upstream genomic sequence. These included transcription factors with established gut function, such as Escargot, Sox100B or Fork head.

**Figure 6.**
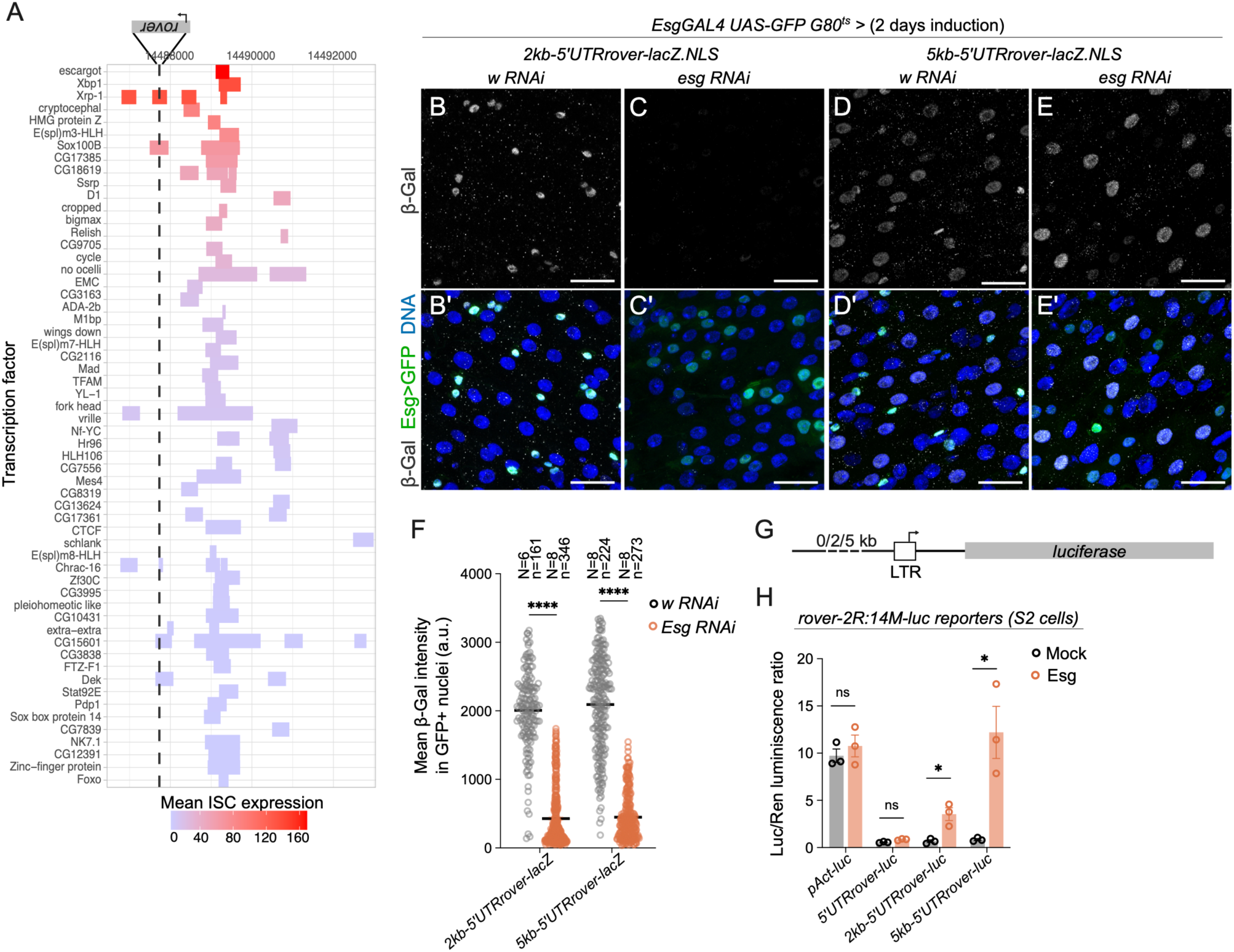
Escargot transcription factor drives *rover-2R:14M* expression *via* the upstream genomic sequence. (**A**) Transcription factor binding sites identified bound to the genomic region upstream of the *rover- 2R:14M-lacZ* locus based on the published ChiP-seq datasets (modENCODE, (67)). Transcription factors were ordered based on mean expression levels in intestinal stem cells (ISCs) obtained from (68). (**B-E’**) Representative confocal images of female midguts with *rover-2R:14M-lacZ* reporter constructs inserted on the X chromosome stained for β-galactosidase (β-gal, grey), GFP (green) and DNA (blue). White RNAi (w, control, B, B’, D, D’) or Esg RNAi (C, C’, E, E’) was induced only in progenitor cells marked with GFP (*EsgGAL4 UAS-GFP G80^ts^*) for two days. Scale bar: 25 μm. (**F**) Quantification of β-galactosidase signal intensities in GFP-positive progenitor cells upon *white* or *esg* knockdown in guts carrying the *rover-2R:14M-lacZ* reporter constructs inserted on the X chromosome. N: number of guts scored; n: number of GFP+ nuceli scored. were applied. **** p<0.0001 (Two-tailed Mann-Whitney test). (**G**) Schematic representation of the *rover-2R:14M* cell culture reporter constructs. Luciferase reporter was placed downstream of the *rover-2R:14M* 5’UTR region replacing all *rover* native open reading frames. 0, 2 or 5 kb of the upstream genomic sequence was also added. (**H**) Luciferase assays for transcriptional activation in S2 cells. Luciferase to renilla luminescence ratios are plotted. *pAct-luc* represents a luciferase reporter under control of a constitutive promotor. All constructs were tested in presence and absence of the Escargot (Esg) transcription factor. ns, non-significant; * p<0.05 (Two-tailed Mann-Whitney tests). The graph is representative of three independent experiments, with three experimental replicates each.

For further analysis we decided to focus on Escargot (Esg), since the observed patterns of *rover-2R:14M- lacZ* reporter activity were reminiscent of *esg* expression, enriched in progenitor cells (ISCs and EBs), and expressed to lower levels in differentiated enterocytes (ECs) (68). In flies carrying the *rover-2R:14M-lacZ* reporters, we performed a transient, progenitor-specific knock-down of Esg using a temperature-sensitive GAL4-UAS system (*Esg>GAL4 UAS-GFP GAL80^ts^*)(69) (***Figure 6B-E’***). Control GFP-marked progenitor cells (expressing *white* RNAi) were positive for β-Gal staining (***Figure 6B, B’, D, D’***), however we observed marked decrease in β-Gal staining in GFP-positive cells upon *esg* knock-down. This effect was observed for both, the *2kb-5’UTRrover-*lacZ (***Figure 6C, C’*** and ***F***) and the *5kb-5’UTRrover-*lacZ (***Figure 6E, E’*** and **F**) reporters, suggesting that Esg could positively regulated *rover* reporter expression *in vivo*.

Importantly, in the fly gut, Esg maintains the progenitor cell fate and its depletion de-regulates many genes, switching on the EC differentiation markers and, in longer term, leading to stem cell loss (70,71). Indeed, visibly enlarged nuclei of GFP-positive cells upon *esg* knock-down indicated ongoing loss of the progenitor fate in these cells. Thus, we cannot exclude that the decrease in the *lacZ* reporter expression was a secondary consequence of deregulation of other genes than Esg itself. Hence, in order to directly test the involvement of Esg in the *rover-2R:14M* transcriptional regulation, we made use of *in vitro* luciferase reporter assays in *Drosophila* S2 cells (***Figure 6G***). Transient transfection of the luciferase reporters under control of the *rover-2R:14M* 5’UTR alone or with addition of the upstream sequences (2 kb or 5 kb) did not result in detectable luciferase activity, as compare to the positive control with a constitutive actin promoter (*pAct-luc*) (***Figure 6H***). However, strikingly, co-expression with Esg led to a significant increase in luciferase activity of *rover-luc* reporters. The transcriptional upregulation was observed only in presence of the genomic sequence (2 kb or 5 kb) upstream of the *rover-2R:14M*, suggesting that the sequence is required for reporter activation and that Escargot acts directly or indirectly on the sequence, regulating reporter expression.

Together these results, obtained with the *in vivo* and cell culture *rover* expression reporters, support the idea that the active *rover-2R:14M* LTR/ERV retrotransposon locus may be regulated by Escargot, a transcription factor critical for cell fate regulation in the fly intestine. This regulation requires the genomic sequence present upstream of the *rover-2R:14M* locus.

## Discussion

Although expression and mobility of retroelements is increasingly reported in diverse organisms and somatic contexts, the regulation of such activity is not well understood. Here, we provide new insight into somatic activity of endogenous LTR/ERV retrotransposons.

Firstly, through whole genome long-read sequencing of *Drosophila* tissues we deliver a detailed landscape of germline and somatic TE insertions in the analyzed genome. As previously reported by us (39), as well as by others (38), our analysis highlights the important and often unappreciated variability in TE composition between fly strains. Indeed, about half of all full-length TE insertions mapped by us are not found in the reference genome. Notably, in contrast to reference TEs, many non-reference TEs are found in euchromatic regions, which, as we show here, may contribute to their transcriptional activity or increase the chance that such insertions influence expression of neighboring genes. This emphasizes the frequently overlooked importance of carefully characterizing or standardizing genetic backgrounds, particularly when somatic TE expression or mobility is studied and compared in different contexts.

Secondly, using our datasets, we address a commonly presumed idea that transposable element activity is unleashed upon aging (52,54). While we report changes in TE transcript levels in the fly gut and head as seen previously (55,56,72,73), we detect no striking increases in *de novo* somatic TE insertions in these tissues. This result, while contradictory with previous, reporter-based data (55,72,74,75), is consistent with recent sequencing-based studies (73,76–78). We cannot exclude that different results could be obtained when systemically analyzing tissues isolated from flies with other genetic backgrounds. Nonetheless, to date, evidence for age-related unleashed mobility of endogenous TEs in the *Drosophila* soma remains scarce. In contrast, we detected many *de novo* retrotransposon insertions in guts isolated from young flies. We cannot eliminate the possibility that these insertions arose during the first few days after eclosion, but the presence of somatic insertions in young adults suggests that retrotransposon mobility could have occurred in pre-adult stages and continued to arise in the adult tissues. This is also consistent with the fact that *rover-2R:14M-lacZ* reporter expression was readily detected in larval as well as adult intestines. Furthermore, the potential developmental TE activity observed by us goes along with recently reported *mdg4* retrotransposon activation during *Drosophila* metamorphosis, proposed to contribute to anti-viral immunity (9).

Further, we deliver new insight into retrotransposon regulation in the fly gut. Through our analysis of *de novo* somatic insertions with long-read DNA sequencing, we identified a “hot” LTR/ERV locus, *rover- 2R:14M.* This example of a somatically mobile LTR/ERV retrotransposon, adds to the previously characterized cases of mobile non-LTR L1 elements in mammalian contexts (26,31,32,35,79). Hence, we were able to investigate copy-specific LTR-element regulation, in line with a growing body of evidence demonstrating that examining TE activity necessitates locus-specific approaches (24,25,80,81), the same way as genes are studied.

To date, studies on the regulation of LTR/ERV elements have been focusing on the sequences carried by the retrotransposons themselves (16–19,40–44). In contrast, we revealed that *rover-2R:14M* is located in permissive chromatin environment, within an intron of an expressed gene, and its transcription is regulated in *cis* by the genomic sequence upstream to the *rover-2R:14M* sequence. Interestingly, in a recent pre- print concomitant with our work, Glaser et al. characterize a polymorphic mouse LTR retrotransposon,

MusD, which achieves its expression during limb development by adopting the expression pattern of neighboring genes contained within the same regulatory domain (82). Thus, this study along with ours, provide complementary examples of how LTR retroelement activity may be regulated tissue-specifically by the genomic environment of the locus. It suggests that such level of regulation is conserved and, as a consequence, there are likely many endogenous retrotransposons regulated in the manner we describe.

In addition to highlighting the role of the genomic environment in the regulation of LTR retroelements, we also identify a transcription factor contributing to this regulation. Numerous transcription factors were shown to bind to retroelement sequences and regulate retrotransposon expression in different biological contexts (83,84). Examples include YY1 (85), RUNX3 (86), p53 (87), SRY (88), MeCP2 (89), SOX2 (90), PAX5 (91) or SOX6 (36). In contrast to these findings, our work implies that regulation of the donor *rover- 2R:14M* locus by a gut lineage-specific transcription factor relies not on the *rover-2R:14M* itself, but on the genomic sequence upstream to the locus. Considering this level of regulation, which has not been investigated so far, the complete spectrum of tissue-specific transcription factors that regulate somatic retrotransposon expression in different biological contexts is certainly yet to be discovered.

Here, we identified Escargot as a regulator of *rover-2R:14M* expression in the fly intestinal tissue. Esg, a snail-family transcription factor, is well-known to control stem cell fate in the fly intestinal lineage (70,71,92,93). In this context, Esg, enriched in progenitor cells, is known to act as a repressor of differentiation genes. Thus, it’s function as a transcriptional activator my appear surprising. However, Esg was shown to bind to a relatively large set of genes, including those with down-regulated expression upon Esg knock-down (70,71). Furthermore, Snail, another closely related transcription factor promoting mesoderm development in the fly embryo through repression of ectodermal genes (94), also functions as a transcriptional activator (95). Thus, it is probable that Esg, perhaps *via* other co-factors and/or transcription factors, may promote transcription of some targets in the fly gut, including of the *rover2R:14M*.

Altogether, our work highlights a new level of locus-specific regulation of LTR/ERV elements by the genomic environment in which these retrotransposons are inserted. This notion is particularly relevant in light of high TE polymorphisms already mentioned above and documented not only in *Drosophila* (48,38,96), but also in human (97–100) or other animal and plant species (101–103). Indeed, the *rover- 2R:14M* locus identified by us is a non-reference polymorphic insertion, as are many of the described somatically active human L1 retrotransposons (35). The full extent to which TE polymorphism contributes to their somatic expression and mobility in different species remains to be addressed.

The biological significance of somatic retrotransposon expression and mobility, although not yet well understood, is now well appreciated and intensively investigated (104). LTR/ERV-type retroelements are not mobile in humans, but they continue to mobilize in other mammalian species. Furthermore, retrotransposon activity may affect host tissues in multiple ways, including transposition-independent, through their transcripts or protein products. Thus, studies as ours, helping to better understand retroelement regulation in diverse lineages are relevant.

## Materials and methods

### Experimental techniques

#### Drosophila stocks

Unless stated otherwise, *Pros>2xGFP (ProsGFP)* stock was used for most sequencing experiments, the same as in (39). This genotype is obtained by crossing ;;*Pros^V1^GAL4/TM6BTbSb* females (J. de Navascués) with *;UAS-2xGFP;* males (Bloomington). The characterized *rover-2R:14M* insertion is present in the ;;*Pros^V1^GAL4/TM6BTbSb* stock, but not in the *;UAS-2xGFP;,* thus it is heterozygous in the *ProsGFP* animals. *w^1118^* stock (gift from M. McVey) was used for RNA-seq as a control stock not carrying the *rover- 2R:14M* insertion. To generate flies carrying the *rover-2R:14M* insertion in an Ago2 mutant background, we isolated the second chromosome carrying *rover-2R:14M* insertion from the ;;*Pros^V1^GAL4/TM6BTbSb* stock and backcrossed the *rover-2R:14M* insertion as well as the *Ago2^414^* mutant allele (from C. Saleh, Institut Pasteur, Paris, France) to the same *w^1118^* background for 7 generations. We then combined the two stocks using standard crosses with balancer lines to obtain flies homozygous for the *rover-2R:14M* insertion and the *Ago2^414^* mutation. *Rover-2R:14M-lacZ* reporter lines were generated in this study (see below). Other stocks used included: UAS*-white RNAi* (BL #33762); *UAS-spn-E RNAi* (BL #34808); *UAS- zuc RNAi* (E. Brasset, iGReD, Clermont-Ferrand, France); *UAS-esg RNAi* (BL #34063); *tj-Gal4* and *nos- Gal4* (L. Teysset, C. Carré, IBPS, Paris, France), *esgGAL4 UAS-GFP GAL80ts* (69). Full genotypes corresponding to main and supplementary figures are listed in the supplementary information file.

#### Fly husbandry

Flies were maintained on a standard medium at 25°C with a day/night light cycle. For crosses, 10–15 females were mixed with males in standard vials. For larval experiments, 3^rd^ instar wandering larvae were selected. Adult progeny was collected over 2–4 days after eclosion and kept at a density of 25-30 flies/tube (mixed sexes). Flies were flipped to fresh tubes every 2-3 days until needed. For aging experiments 3 time-points were analyzed: young 5-7-day-old, mid-age 20-25-days-old and aged 50-60-days-old. For temporal induction of *white* or *esg* RNAi in adult gut progenitor cells (using *esgGAL4 UAS-GFP GAL80ts* (69)), crosses were maintained in 18°C and 5-10-days-old adult flies were switched to 29°C for 2 days. Females were used for most experiments, unless stated otherwise.

#### Genomic DNA isolation

Tissues were dissected in ice-cold, nuclease-free PBS and snap-frozen in liquid nitrogen before DNA isolation. High molecular weight genomic DNA was isolated from pools of 60 guts or 60 heads with the MagAttract HMW DNA Kit (Qiagen #67563) or *Quick*-DNA HMW MagBead Kit (Zymo Research #D6060) according to manufacturers’ instructions, with tissue lysis performed overnight at 55°C. gDNA was eluted with nuclease-free water. DNA integrity was verified on a 0.6% agarose gel and concentrations were measured with Qubit dsDNA Broad Range Assay Kit. All samples had A260/280 ratios above 1.8 and A260/230 ratios above 2.0.

#### DNA sequencing

Whole genome long-read DNA sequencing libraries were prepared with 500-800ng of DNA following the 1D Genomic DNA by Ligation Protocol (SQK-LSK109, Oxford Nanopore Technologies). Sequencing was performed on MinION or GridION using R9.4.1 flow cells (Oxford Nanopore Technologies) and 48 h-long sequencing runs. Supplementary Table 1 provides basic sequencing statistics for all samples.

#### RNA isolation

For RNA isolation, gut and head tissues were dissected in cold, RNase-free PBS, transferred to 100 μl of TRIzol Reagent (Thermo Fisher Scientific), homogenized with a plastic pestle and snap-frozen in liquid nitrogen for storage at −80°C. Upon thawing, samples were further processed according to the TRIzol Reagent manufacturer’s protocol. Purified RNA was treated with DNase (Ambion) for 1 h at 37°C, further purified with phenol-chloroform extraction and isopropanol precipitation, and resuspended in RNase-free water. All samples had A260/280 ratios above 1.9 and A260/230 ratios above 2.0. RNA integrity was checked on Bioanalyzer (Agilent) using the Agilent RNA 6000 Nano Kit and concentrations were assayed with the Qubit RNA Broad Range Assay Kit (Thermo Fisher Scientific).

#### RNA sequencing

For the short-read transcriptome analysis, 700ng of total RNA was used to prepare libraries according to the TruSeq Stranded mRNA protocol (Illumina). Samples were processed in biological triplicates. 2X100 bp paired-end sequencing was performed on Novaseq (Illumina). For long-read Oxford Nanopore (ONT) cDNA sequencing, we first prepared mRNA with Dynabeads mRNA Purification Kit (Invitrogen #61006) starting from 100 μg of DN-ase digested total RNA, according to the manufacturer’s protocol. Supplementary Table 1 provides basic sequencing statistics for all samples. mRNA concentration was quantified with the Qubit RNA Broad Range Assay Kit (Thermo Fisher Scientific) and sample quality was checked on Bioanalyzer (Agilent). Samples were then prepared according to the protocol for ONT Direct cDNA sequencing (SQK-DCS109, Oxford Nanopre), using 150 ng of purified mRNA as input. Samples were run on MinION using R9.4.1 flow cells (Oxford Nanopore Technologies) and 48 h-long sequencing runs.

### Generation of *lacZ* and *luciferase* reporters

To construct reporter plasmids, we amplified the *rover-2R:14M* 5’UTR region alone or with its upstream genomic sequence of 2 or 5 kb from *ProsGFP* genomic DNA. These sequences were then cloned into appropriate vectors using NEBuilder® HiFi DNA Assembly protocol (New Englad Biolabs #E5520). lacZ reporter plasmids were obtained by replacing the *hsp70* promoter sequence from the placZ-attB (105) vector upstream of the *lacZ* gene with the regulatory sequences of interest. The plasmids were then injected by Bestgene (Chino Hills, CA, USA) for integration into two different landing sites: VK38 (Ch X, Bloomington stock #9753) and VK05 (Ch 3L, Bloomington stock #9725). For S2 cell reporter plasmids, we used the same cloning strategy to place the *rover-2R:14M* regulatory sequences into the pAct-GL3 vector (Promega) upstream of the luciferase gene. pAct-Renilla vector was used to correct for transfection efficiency. All final constructs were checked by ONT whole plasmid sequencing (Eurofins Genomics).

#### Immunolabeling

Midguts were dissected and fixed in 4 % PFA (FISHER SCIENTIFIC S.A.S. 15828264), 1X PBS for 3 hours. Fixed tissues were washed with PBT (1X PBT, 0.1% Triton X-100) and incubated for at least 30 minutes in 50% Glycerol, 1X PBS, followed by placing back into PBT, to allow the waste to exit by osmotic pressure. The tissues were then incubated with primary antibodies at 4°C overnight, washed three times with PBT (20 min each) and incubated with the secondary antibodies for 3 hr at room temperature. Next, the guts were washed three times in PBT and stained with DAPI at the last wash. Stained tissues were equilibrated in 50% Glycerol, 1X PBS for at least 1 hour before mounting on microscopy slides in mounting medium. Ovaries were dissected in 1X PBS, fixed in 4% PFA for 20 minutes, and washed 3 x 20 minutes in PBT (PBS, 0.3% Triton X-100). Ovaries were then incubated in a blocking solution (PBTA: PBT, 2% BSA – Sigma A3059) for a minimum of 30 minutes. Primary antibodies were diluted in PBTA and ovaries were incubated in this solution overnight at 4°C. Tissues were then washed in PBT 3x10min, and incubated in PBTA for at least 30 minutes before incubation in secondary antibodies in PBTA for 3 hours at room temperature. Finally, tissues were washed 3x10 min in PBT, and mounted in DABCO (D27602, Sigma) with 70% glycerol.

The following primary antibodies were used: chicken anti-β-Gal (Abcam #9361), mouse anti-Prospero (DSHB #MR1A-c).

#### Microscopy

Images were obtained using an upright confocal laser scanning microscope TCS SP8, Leica (Leica Microsystems, Germany) using a HC PL APO CS2 63x/1.4 oil immersion objective lens. 12 bit numerical images were acquired with the Leica Application Suite X software (LAS Version 3.5.6; Leica, Germany) and processed using Fiji (ImageJ, (106)) version 1.53c. Adult and larval guts were imaged in the posterior R4 region (according to (107)). To quantify β-Gal signal (Figure 6F), we used a custom macro available upon request. Briefly, a maximum intensity projection of the Z-stacks was generated and a binary mask was created on the green channel. Then, Analyze Particles function was used to delineate individual cells (criteria size, larger than 10 µm²) and mean fluorescence intensities were measured for each cell, with measurements restricted to the defined regions of interest (ROIs) corresponding to the GFP positive cells.

#### S2 cell culture

Drosophila Schneider’s S2 cells (DGRC #181) were maintained at 25°C in Schneider’s Drosophila medium (Invitrogen) supplemented with 10% heat inactivated fetal calf serum (Gibco) and antibiotics (penicillin/streptomycin, Invitrogen). Cells in the exponential phase of growth were used for all the experiments.

#### Luciferase assays

S2 cells were transiently transfected with Effectene Transfection Reagent (Qiagen # 301425) according to the manufacturer’s protocols, to deliver the following vector: 1- luciferase reporter plasmid, 2- TF overexpression plasmid under constitutive promoter (pAc5.1), and 3- a Renilla construct for normalization. The ratio of luciferase:renilla plasmids was kept at 10:1. The total amount of transfected DNA was kept constant by adjusting with an empty vector. Levels of Luciferase and Renilla were measured 48 hours after transfection using Dual-Glo Luciferase Assay System (Promega #E2920) and Spectramaax M5 (Molecular Devices). For each transfection, at least three biological replicates were performed, each done in triplicate. cDNA for Esg was amplified from the DGRC Stock number 1645028 (RRID:DGRC_1645028).

### Computational analysis

#### Processing of long-read DNA-seq samples

Raw sequencing reads from the ONT DNA sequencing libraries generated in this study and in the ONT DNA sequencing libraries from (39) were basecalled using guppy v6.0.1, a basecaller developed by ONT, with the dna_r9.4.1_450bps_hac model. Basecalled reads were merged into a single FASTQ file for each library. Sequencing adapters were trimmed using Porechop v0.2.4 (https://github.com/rrwick/Porechop). The reads were filtered using NanoFilt v2.8.0 (108) with filters for a minimum average read quality score (-q 10) and a minimum read length (-l 500). The reads were aligned to the FlyBase dm6.48 (109) reference genome using minimap2 (110) with -x map-ont preset and the parameter for to retain the MD tag (--MD). The resulting alignments were sorted, filtered for a minimum mapping quality 5, and indexed using samtools v1.13 (111). Sequencing and alignment quality were assessed with NanoPlot v1.36.2 (112) and pycoQC v2.5.2 (113).

#### Calling non-reference TE insertions in long-read DNA-seq samples

tldr v1.2.2 (81) was used to call non-reference insertions with the following parameters: --color_consensus --trdcol --detail_output --minreads 1 --min_te_len 500 --max_cluster_size 100. The TE consensus sequences for *D. melanogaster* species made available by the Bergman’s lab (https://github.com/bergmanlab/drosophila-transposons) were used as the reference TE library. The following criteria were used to filter raw calls:

1. the fraction of the inserted sequence covered by the TE sequence ≥ 0.5;
2. the length of the inserted sequence ≥ 500bp;
3. the median mapping quality score of the reads supporting the insertion ≥ 30;
4. the chromosome of the insertion was the autosomes or chrX;
5. the coverage at the integration site ≥ 10.

Calls meeting the following criterion were classified as full-length insertions: *(Total length of the insertion)*

- *(Fraction of the inserted sequence covered by the TE sequence)* / *(Consensus TE length)* ≥ 0.7. All other calls were classified as truncated insertions. Next, calls from all samples were clustered based on subfamily, genomic breakpoint coordinates (allowing 100bp margin) and DNA strand. Clusters were reviewed and filtered based on the presence of full-length calls, TSDs, the minimum coverage at breakpoints, status of the ‘remappable’ filter in the tldr output. Clusters were retained if they satisfied the following conditions:

- at least one call in the cluster was for a full-length insertion;
- the breakpoint region had coverage < 200;
- coverage at the integration site in at least one sample was ≥ 15;
- at least one call in the cluster passed ‘remappable’ filter in the tldr ouput. The calls in each retained cluster were collapsed, and
- left genomic breakpoint of the cluster was defined as the median of all left genomic breakpoints in the cluster;
- right genomic breakpoint of the cluster was defined as the median of all right genomic breakpoints in the cluster;
- ONT read ratio of the cluster was defined as the median of all ONT read ratios in the cluster;
- TSDs as well as samples and tissues that supported each cluster were recorded.

Clusters within 1kb of a reference insertion of the same sub-family were removed, as they likely represented misalignments or DNA base events. This was supported by the fact that most such clusters included calls with TSDs > 100bp and/or long non-TE flanking regions.

The remaining clusters were further iterated over, and clusters within a 2kb margin of genomic breakpoints for the same sub-family were extracted. The following rules were applied:

1. if no cluster in the subsection had the ‘PASS’ filter flag in the tldr output, only the cluster with the highest number of supporting samples was kept;
2. if only one cluster in the subsection had the ‘PASS’ filter flag in the tldr output, only this cluster was kept;
3. if multiple clusters in the subsection had the ‘PASS’ filter flag in the tldr output, the cluster supported by both tissues was kept (if such a cluster was present in the subsection); otherwise, all clusters in the subsection were kept.

#### Calling reference TE insertions

RepeatMasker v4.1.2 (Smit, AFA, Hubley, R & Green, P. RepeatMasker Open-4.0) with CONS- Dfam_withRBRM_3.5 library (114) was used to mask dm6.48 reference genome. one_code_to_find_them_all v1.0 (115) was used to assemble the masked sequences into complete TE copies. Insertions that were longer than 70% of the respective consensus sequences and with less than 30% sequence divergence relative to the respective consensus sequences were called full-length. To detect reference insertions present in the genome of *ProsGFP* strain, reads with primary alignments with mapping quality > 5 spanning regions upstream and downstream of a reference insertion were extracted from the ONT bam files using custom Python code and the pysam module (https://github.com/pysam-developers/pysam). Reads with alignments of at least 100bp within reference TE coordinates were classified as supporting the insertion, while all other reads were classified as opposing the insertion. A reference insertion was considered present in the genome if it was supported by at least two reads in at least two ONT samples.

#### Genotyping TE calls in ONT samples

The clustered non-reference and reference calls were genotyped in two steps. Initial genotype assignments were made based on the following criteria:

- non-reference calls supported by a single read and detected only in one ONT sample were assigned ‘singleton’ genotype;
- non-reference calls with ONT read ratio < 0.1 that were detected in both tissues were assigned ‘rare’ genotype;
- non-reference calls with ONT read ratio ≥ 0.1 were assigned ‘fixed’ genotype;
- reference calls were assigned ‘reference’ genotype;
- all other calls were considered as ‘ungenotyped’.

Calls detected in only one tissue were further examined as follows. For each such call, the clipped parts of reads longer than 100bp were extracted from all samples, except the sample where the call was detected, within a 200bp margin of the left genomic breakpoint, using pysam Python module. The extracted parts were mapped to the consensus TE sequences with mappy Python module. Primary alignments with a mapping quality ≥ 15 were retained. The genotype, sample and tissue supporting a call were updated if an alignment hit was found for the same subfamily. This procedure updated approximately 5% singleton calls, 30% of ungenotyped calls, and 3% of fixed non-reference calls. The updated calls were supplemented with the Illumina VAF, the number of Illumina sample pairs supporting a call, and the genotype inferred from the Illumina samples if a call matched a call in the Illumina samples for the same subfamily, on the same chromosome, and within 100bp margin of the genomic breakpoints.

Among the ungenotyped calls, 73 calls with supporting reads that poorly mapped to one TE consensus sequence and 4 calls with supporting reads that mapped to several TE consensus sequences were removed. To archieve this, mapping quality, the number of reads, and TE subfamily information for each call were obtained from the tldr detail.out output files. Among the remaining ungenotyped calls:

- 117 insertions were singletons without a TSD;
- 88 insertions were supported by an ambiguous set of samples (e.g., two samples from the same tissue);
- 87 insertions could potentially be considered somatic insertions of high clonality (i.e., insertions detected in one sample but supported by more than one read);
- 56 insertions were singletons from germline-active sub-families.
- 12 insertions were singletons with a TSD longer than 50bp.

#### Calling non-reference germline TE insertions in short-read DNA-seq samples

The coordinates for somatic insertions detected in the short-read Illumina samples from (39) were obtained from Supplementary Table 2. The coordinates for the germline insertions detected in the short-read Illumina samples from (39) were obtained from Supplementary Table 7. The same Illumina samples (samples P7-P66, accession number PRJNA641572) were reanalyzed using a second independent bioinformatics approach. For this, ngs_te_mapper2 v1.0.2 (116) was used with default parameters, except for --min_af 0.01. Calls from ngs_te_mapper2 in all samples were collapsed based on subfamily, breakpoint coordinates and strand, allowing a 50bp margin for the coordinates. The resulting callset was merged with the somatic and germline callsets from (39), mentioned above, based on subfamily and breakpoints coordinates, also allowing 50bp margin for the coordinates. The samples and the tissue supporting each insertion were recorded. The VAF of an insertion was defined as the average of the VAFs in the supporting samples. An insertion was assigned a genotype based on the following criteria:

- ‘germline’ if it was detected in at least 3 pairs of samples (i.e. gut and head samples from one fly);
- ‘private germline’ if it was detected in exactly one sample pair;
- ‘rare germline’ if it was detected in > 1 and < 3 sample pairs.

#### Definition of germline active sub-families

The sub-families were sorted in descending order based on the total sum of ‘rare germline’ and ‘private germline’ insertions detected in the Illumina samples. A cutoff of 5 insertions was set to define the germline-active sub-families.

#### Normalization of raw singleton counts

To normalize raw singleton counts in a sample, the raw singleton count of a sub-family was multiplied by 1000 and divided by the number of reads longer than 3.5 times the respective consensus TE length in the sample. This approach normalized raw counts for sequencing depth, considering only reads whose both flanks could potentially be aligned to the genome in the presence of a TE insertion, as this is a necessary condition for detecting a singleton insertion. The number of reads in a sample was calculated by extracting the reads with primary alignments from the respective BAM file and calculating their length.

A one-way ANOVA test and Holm’s multiple test correction method were used to compare the mean normalized counts between three age groups.

#### Creating consensus sequences for *rover* fixed reference and non-reference insertions

To create consensus sequences for reference insertions, the parts of the reads between reference TE coordinates that had alignments longer than 200bp within the coordinates were extracted from the ONT BAM files with custom Python code. Multiple sequence alignments of the extracted sequences supporting each reference insertion were performed using MAFFT (117), and the consensus sequences were created using the cons tool from the EMBOSS package (118).

To create consensus sequences for non-reference fixed insertions, the ONT BAM files were parsed with custom Python code, and inserted sequences (‘I’ in CIGAR string or ‘1’ in CIGAR tuple in pysam) between genomic breakpoint coordinates were extracted. Extracted sequences supporting an insertion were aligned to the *rover* consensus sequence with minimap2. Alignments were sorted and indexed with samtools view and samtools index. A consensus sequence for an insertion was created with samtools consensus.

#### Identification of sequence variants supporting *rover* donor locus

Singletons in each ONT sample were aligned to the *rover* consensus sequence using minimap2, sorted and indexed using samtools, and visually inspected in IGV (119).

The consensus sequences of 15 reference and 5 fixed non-reference *rover* insertions present in the genome of the *ProsGFP* strain were first visually compared to *rover* consensus sequence by generating and visualizing multiple sequence alignments with Mauve (120). Insertions with large structural variants (SVs), that were not present in singletons were excluded from the subsequent analysis. The sequences of the remaining 4 reference and 4 fixed non-reference insertions were aligned to the *rover* consensus sequence using minimap2, sorted and indexed with samtools, and visually inspected in IGV. Visual inspection of 50-300bp long SVs present in singletons and fixed insertions allowed to rule out additional 6 fixed insertions. The *rover-2R:14M* and *rover-2L:18M* insertions differed only by a set of 10 SNVs. The fraction of singletons supporting each SNV present in the *rover-2R:14M* and not present in the *rover- 2L:18M* was quantified using freebayes https://doi.org/10.48550/arXiv.1207.3907

#### Creating genome and transcriptome references for expression analysis

To quantify *rover* consensus expression, the masked dm6.48 genome was supplemented with an artificial chrTE that consisted of consensus sequences of all TEs present in the *D. melanogaster* TE library. To quantify *rover* per copy expression, the *rover* sequence in chrTE was masked, and an additional chrRS was added, that consisted of sequences of all fixed *rover* insertions present in the genome of the *ProsGFP* strain.

#### Illumina RNA sequencing analysis

To quantify *rover* consensus expression, Illumina RNA-seq libraries from (39) and Illumina RNA-seq libraries generated in this study were aligned to the masked reference genome plus chrTE using STAR 2.7.10a (121) with the following parameters: --sjdbOverhang 100 --outFilterMultimapNmax 1 -- winAnchorMultimapNmax 10 --outMultimapperOrder Random --outFilterMismatchNoverLmax 0.3 -- quantMode GeneCounts quantification mode. The expression count tables were grouped for all samples, and the counts were normalized using TMM normalization method from the edgeR (122) R package. Differential expression analysis was performed using DESeq2 (123) R package, and the volcano plots were plotted using the ggplot2 package (H. Wickham. ggplot2: Elegant Graphics for Data Analysis. Springer-Verlag New York, 2016).

To quantify *rover* per copy expression, FASTQ files with the reads that had primary alignments within the *rover* consensus sequence region on the chrTE in the BAM files produced by STAR were created using view, collate, and fastq from samtools. These were then realigned to the masked chrTE plus chrRS genome using STAR 2.7.10a with the following parameters: --outFilterMultimapNmax 57 -- winAnchorMultimapNmax 100 --outFilterMismatchNmax 999 --outFilterMismatchNoverLmax 0.01 -- quantMode GeneCounts quantification mode. The expression count tables were grouped for all samples. A new count table was created that included gene and consensus TE counts (excluding *rover*) from the mapping to the masked reference genome plus chrTE, and *rover* fixed insertions counts from the mapping to masked chrTE plus chrRS. The counts were normalized using TMM normalization method from the edgeR R package.

#### Analysis of ONT cDNA sample

Raw reads were basecalled with guppy v6.0.1 using the following parameters for the flowcell and sequencing kit: --flowcell FLO-MIN106 --kit SQK-DCS109. Basecalled reads were trimmed and oriented with pychopper v2, developed by ONT.

To quantify *rover* consensus expression, basecalled reads were aligned to the masked reference genome plus chrTE using minimap2 with the -x splice preset and the --secondary=no parameter. Read counts were quantified using featureCounts v2.0.1 from the Subread package (124) with the following parameters: -Q 5 -L. Raw read counts are reported in Figure 3.

To quantify *rover* per copy expression, FASTQ files with the reads that had primary alignments within the *rover* consensus sequence region on the chrTE in the BAM files produced by STAR were created using view, collate, and fastq from samtools. These were then realigned to the masked chrTE plus chrRS genome using minimap2 with the -x map-ont preset and the --secondary=no parameter. Reads were quantified with featureCounts as described above.

#### Motif analysis of the LTR region

Position weight matrices for the components of the PolII transcription initiation complex (TATA-box, Inr, DPE and MTE) were created from the frequency tables from (125) using TFBSTools (126) in R. The first 363b of the *rover-2R:14M*, *rover-2L:18M,* and *rover-2R:21M* insertions were scanned with parameters for a minimum motif score 80% and the ‘+’ strand using the searchSeq function in TFBSTools. Motif scores were scaled to 1 separately for each element and plotted using custom Python scripts.

#### Motif analysis of the internal 2kb region

Position weight matrices from the JASPAR database (127) for transcription factor binding profiles were used to scan the internal 2kb regions of the *rover-2R:14M*, *rover-2L:18M* and *rover-2R:21M* insertions with parameters for a minimum motif score 80% and the ‘+/-’ strands, in the same way as described above. Motif scores were scaled to 1 separately for each transcription factor. Expression values for each transcription factor in ISC, EB, EC, EE cell types were taken from (68).

### Motif analysis of the *rover-2R:14M* upstream region

ChIP-seq BED tracks with peaks for the *D. melanogaster* dm6 reference genome were downloaded from modENCODE (67). The tracks were intersected with the upstream genomic region for the *rover-2R:14M* locus, and transcription factors expressed in ISC, based on the expression data from (68), were plotted with custom R scripts.

#### Epigenetic data analysis

Heatmaps for DamID tracks for PolII, Brm, Pc, HP1, H1 factors for ISC, EB, EC, EE and ATAC-seq tracks for ISC from (63) (accession number PRJNA933194) were plotted for 10kb upstream genomic regions of the full-length fixed *rover* insertions using computeMatrix and plotHeatmap from deepTools (128).

## Data and code availability

Sequencing data generated for this study have been deposited at GEO (BioProject ID PRJNA1202082; ONT DNA-seq: PRJNA1202082, RNA-seq and ONT cDNA-seq: GSE285324) and are publicly available. Accession numbers of datasets published previously are provided in the Supplementary Table 1. All original code is available from https://github.com/nrubanova/rover_ERV.

## Materials availability

All reagents generated in this study, including plasmids and *Drosophila* stocks, are available from the lead contact upon request.

## Supporting information

Rubanova etal-Supplementary File

Rubanova etal-Supplementary Table 1

Rubanova etal-Supplementary Table 2

Rubanova etal-Supplementary Table 3

## Acknowledgements

We thank Marius van den Beek, who’s initial ONT DNA-seq data analysis led to the identification of the *rover-2R:*14M locus. We would also like to acknowledge S. Chambeyron for sharing and discussing unpublished data, A. Boivin for his comments on the manuscript, as well as J. Crocker, L. Teysset, C. Carré, E. Brasset, C. Saleh, Bloomington, and the Drosophila Genomics Resource Center (NIH Grant 2P40OD010949) for providing fly stocks or reagents. The high-throughput sequencing involved in this study was performed by the ICGex NGS platform of the Institut Curie (supported by the grants ANR- 10-EQPX-03 (Equipex) and ANR-10-INBS-09-08 (France Génomique Consortium) from the Agence Nationale de la Recherche (“Investissements d’Avenir” program), by the Canceropole Ile-de-France, and by the SiRIC-Curie program—SiRIC Grant “INCa-DGOS4654), as well as the I2BC High-throughput sequencing facility, supported by France Génomique (funded by the French National Program “Investissement d’Avenir” ANR-10-INBS-09). The present work has also benefited from Imagerie-Gif core facility supported by I’Agence Nationale de la Recherche (ANR-10-INBS- 04/FranceBioImaging; ANR-11- IDEX-0003-02/Saclay Plant Sciences). We thank Valerie Nicolas for her assistance in quantification of confocal images. This work was supported by the Fondation pour la Recherche Médicale (A.J.B., EQ202003010251) and the ERC StG gutTEimpact (K.S., 101078070). Salary support of KS is from Inserm; AJB and MP from CNRS, FC from University Paris-Saclay, SN from University of Versailles St-Quentin and LB from Ministère de l’Enseignement Supérieur (doctoral grant).

## Author contributions

N.R., A.J.B. and K.S. designed the study. N.S., A.J.B. and K.S. supervised the work. N.R. performed all bioinformatic analysis with a contribution of L.B. to Fig 6A. D.S, L.B., F.C., S.N. and M.P. and K.S. performed experiments. A.J.B. and K.S. provided funding. N.R. and K.S. wrote the manuscript with contribution of A.J.B. and all other authors.

## Declaration of interests

Authors declare no conflicts of interests.

## Notes

### Competing Interest Statement

The authors have declared no competing interest.

## References

1. Wells JN, Feschotte C. A Field Guide to Eukaryotic Transposable Elements. Annu Rev Genet. 2020 Nov 23;54(1):539–61.

2. Reus K, Mayer J, Sauter M, Zischler H, Müller-Lantzsch N, Meese E. HERV-K(OLD): Ancestor Sequences of the Human Endogenous Retrovirus Family HERV-K(HML-2). J Virol. 2001 Oct;75(19):8917–26.

3. Cosby RL, Chang NC, Feschotte C. Host–transposon interactions: conflict, cooperation, and cooption. Genes Dev. 2019 Sep 1;33(17–18):1098–116.

4. Dopkins N, O’Mara MM, Lawrence E, Fei T, Sandoval-Motta S, Nixon DF, et al. A field guide to endogenous retrovirus regulatory networks. Molecular Cell. 2022 Oct 20;82(20):3763–8.

5. Jachowicz JW, Bing X, Pontabry J, Bošković A, Rando OJ, Torres-Padilla ME. LINE-1 activation after fertilization regulates global chromatin accessibility in the early mouse embryo. Nat Genet. 2017 Oct;49(10):1502–10.

6. Chang NC, Wells JN, Wang AY, Schofield P, Huang YC, Truong VH, et al. Gag proteins encoded by endogenous retroviruses are required for zebrafish development. bioRxiv. 2024 Jan 1;2024.03.25.586437.

7. Simon M, Meter MV, Ablaeva J, Ke Z, Gonzalez RS, Taguchi T, et al. LINE1 Derepression in Aged Wild-Type and SIRT6-Deficient Mice Drives Inflammation. Cell Metabolism. 2019 Apr 2;29(4):871–885.e5.

8. Zhao Y, Oreskovic E, Zhang Q, Lu Q, Gilman A, Lin YS, et al. Transposon-triggered innate immune response confers cancer resistance to the blind mole rat. Nat Immunol. 2021 Oct;22(10):1219–30.

9. Wang L, Tracy L, Su W, Yang F, Feng Y, Silverman N, et al. Retrotransposon activation during Drosophila metamorphosis conditions adult antiviral responses. Nat Genet. 2022 Dec;54(12):1933– 45.

10. Mietz JA, Fewell JW, Kuff EL. Selective activation of a discrete family of endogenous proviral elements in normal BALB/c lymphocytes. Mol Cell Biol. 1992 Jan;12(1):220–8.

11. Deininger P, Morales ME, White TB, Baddoo M, Hedges DJ, Servant G, et al. A comprehensive approach to expression of L1 loci. Nucleic Acids Res. 2017 Mar 17;45(5):e31–e31.

12. Pehrsson EC, Choudhary MNK, Sundaram V, Wang T. The epigenomic landscape of transposable elements across normal human development and anatomy. Nat Commun. 2019 Dec 10;10(1):1–16.

13. Chung N, Jonaid GM, Quinton S, Ross A, Sexton CE, Alberto A, et al. Transcriptome analyses of tumor-adjacent somatic tissues reveal genes co-expressed with transposable elements. Mobile DNA. 2019 Dec;10(1):39.

14. Ansaloni F, Scarpato M, Di Schiavi E, Gustincich S, Sanges R. Exploratory analysis of transposable elements expression in the C. elegans early embryo. BMC Bioinformatics. 2019 Nov;20(S9):484.

15. Treiber CD, Waddell S. Transposon expression in the Drosophila brain is driven by neighboring genes and diversifies the neural transcriptome. Genome Res [Internet]. 2020 Sep 24 [cited 2021 Feb 9]; Available from: http://genome.cshlp.org/content/early/2020/10/14/gr.259200.119

16. Burn A, Roy F, Freeman M, Coffin JM. Widespread expression of the ancient HERV-K (HML-2) provirus group in normal human tissues. PLOS Biology. 2022 Oct 18;20(10):e3001826.

17. Carter TA, Singh M, Dumbović G, Chobirko JD, Rinn JL, Feschotte C. Mosaic cis-regulatory evolution drives transcriptional partitioning of HERVH endogenous retrovirus in the human embryo. eLife. 2022 Feb 18;11:e76257.

18. She J, Du M, Xu Z, Jin Y, Li Y, Zhang D, et al. The landscape of hervRNAs transcribed from human endogenous retroviruses across human body sites. Genome Biol. 2022 Nov 3;23(1):231.

19. Chang NC, Rovira Q, Wells J, Feschotte C, Vaquerizas JM. Zebrafish transposable elements show extensive diversification in age, genomic distribution, and developmental expression. Genome Res. 2022 Jul;32(7):1408–23.

20. Faulkner GJ, Garcia-Perez JL. L1 Mosaicism in Mammals: Extent, Effects, and Evolution. Trends in Genetics. 2017 Nov;33(11):802–16.

21. Lanciano S, Cristofari G. Measuring and interpreting transposable element expression. Nature Reviews Genetics. 2020 Jun 23;1–16.

22. Athanikar JN, Badge RM, Moran JV. A YY1-binding site is required for accurate human LINE-1 transcription initiation. Nucleic Acids Research. 2004 Jul 1;32(13):3846–55.

23. Lavie L, Maldener E, Brouha B, Meese EU, Mayer J. The human L1 promoter: Variable transcription initiation sites and a major impact of upstream flanking sequence on promoter activity. Genome Res. 2004 Nov 1;14(11):2253–60.

24. Philippe C, Vargas-Landin DB, Doucet AJ, van Essen D, Vera-Otarola J, Kuciak M, et al. Activation of individual L1 retrotransposon instances is restricted to cell-type dependent permissive loci. Burns K, editor. eLife. 2016 Mar 26;5:e13926.

25. Berrens RV, Yang A, Laumer CE, Lun ATL, Bieberich F, Law CT, et al. Locus-specific expression of transposable elements in single cells with CELLO-seq. Nat Biotechnol. 2022 Apr;40(4):546–54.

26. Gerdes P, Chan D, Lundberg M, Sanchez-Luque FJ, Bodea GO, Ewing AD, et al. Locus-resolution analysis of L1 regulation and retrotransposition potential in mouse embryonic development. Genome Res. 2023 Sep;33(9):1465–81.

27. Baillie JK, Barnett MW, Upton KR, Gerhardt DJ, Richmond TA, De Sapio F, et al. Somatic retrotransposition alters the genetic landscape of the human brain. Nature. 2011 Nov;479(7374):534– 7.

28. Evrony GD, Cai X, Lee E, Hills LB, Elhosary PC, Lehmann HS, et al. Single-Neuron Sequencing Analysis of L1 Retrotransposition and Somatic Mutation in the Human Brain. Cell. 2012 Oct;151(3):483–96.

29. Evrony GD, Lee E, Mehta BK, Benjamini Y, Johnson RM, Cai X, et al. Cell Lineage Analysis in Human Brain Using Endogenous Retroelements. Neuron. 2015 Jan 7;85(1):49–59.

30. Upton KR, Gerhardt DJ, Jesuadian JS, Richardson SR, Sánchez-Luque FJ, Bodea GO, et al. Ubiquitous L1 Mosaicism in Hippocampal Neurons. Cell. 2015 Apr;161(2):228–39.

31. Tubio JMC, Li Y, Ju YS, Martincorena I, Cooke SL, Tojo M, et al. Extensive transduction of nonrepetitive DNA mediated by L1 retrotransposition in cancer genomes. Science. 2014 Aug 1;345(6196):1251343–1251343.

32. Scott EC, Gardner EJ, Masood A, Chuang NT, Vertino PM, Devine SE. A hot L1 retrotransposon evades somatic repression and initiates human colorectal cancer. Genome Res. 2016 Jun;26(6):745– 55.

33. Rodriguez-Martin B, Alvarez EG, Baez-Ortega A, Zamora J, Supek F, Demeulemeester J, et al. Pan-cancer analysis of whole genomes identifies driver rearrangements promoted by LINE-1 retrotransposition. Nat Genet. 2020 Mar;52(3):306–19.

34. Nam CH, Youk J, Kim JY, Lim J, Park JW, Oh SA, et al. Widespread somatic L1 retrotransposition in normal colorectal epithelium. Nature [Internet]. 2023 May 10 [cited 2023 May 11]; Available from: https://www.nature.com/articles/s41586-023-06046-z

35. Sanchez-Luque FJ, Kempen MJHC, Gerdes P, Vargas-Landin DB, Richardson SR, Troskie RL, et al. LINE-1 Evasion of Epigenetic Repression in Humans. Molecular Cell. 2019 Aug 8;75(3):590–604.e12.

36. Bodea GO, Botto JM, Ferreiro ME, Sanchez-Luque FJ, De Los Rios Barreda J, Rasmussen J, et al. LINE-1 retrotransposons contribute to mouse PV interneuron development. Nat Neurosci. 2024 Jul;27(7):1274–84.

37. Gagnier L, Belancio VP, Mager DL. Mouse germ line mutations due to retrotransposon insertions. Mobile DNA. 2019 Apr 13;10(1):15.

38. Mérel V, Boulesteix M, Fablet M, Vieira C. Transposable elements in Drosophila. Mobile DNA. 2020 Dec;11(1):23.

39. Siudeja K, van den Beek M, Riddiford N, Boumard B, Wurmser A, Stefanutti M, et al. Unraveling the features of somatic transposition in the Drosophila intestine. The EMBO Journal. 2021 Feb 26;n/a(n/a):e106388.

40. Araujo P, Casacuberta J, Costa A, Hashimoto R, Grandbastien MA, Van Sluys MA. Retrolyc1 subfamilies defined by different U3 LTR regulatory regions in the Lycopersicon genus. Mol Gen Genomics. 2001 Sep;266(1):35–41.

41. Mugnier N, Biémont C, Vieira C. New Regulatory Regions of Drosophila 412 Retrotransposable Element Generated by Recombination. Molecular Biology and Evolution. 2005 Mar;22(3):747–57.

42. Göke J, Lu X, Chan YS, Ng HH, Ly LH, Sachs F, et al. Dynamic Transcription of Distinct Classes of Endogenous Retroviral Elements Marks Specific Populations of Early Human Embryonic Cells. Cell Stem Cell. 2015 Feb;16(2):135–41.

43. Ito J, Sugimoto R, Nakaoka H, Yamada S, Kimura T, Hayano T, et al. Systematic identification and characterization of regulatory elements derived from human endogenous retroviruses. PLOS Genetics. 2017 Jul 12;13(7):e1006883.

44. Geng LN, Yao Z, Snider L, Fong AP, Cech JN, Young JM, et al. DUX4 Activates Germline Genes, Retroelements, and Immune Mediators: Implications for Facioscapulohumeral Dystrophy. Developmental Cell. 2012 Jan;22(1):38–51.

45. Malik HS, Henikoff S, Eickbush TH. Poised for Contagion: Evolutionary Origins of the Infectious Abilities of Invertebrate Retroviruses. Genome Res. 2000 Sep 1;10(9):1307–18.

46. Terzian C, Pélisson A, Bucheton A. Evolution and phylogeny of insect endogenous retroviruses. BMC Evol Biol. 2001 Aug 10;1(1):3.

47. Riddiford N, Siudeja K, Beek M van den, Boumard B, Bardin AJ. Evolution and genomic signatures of spontaneous somatic mutation in Drosophila intestinal stem cells. Genome Res. 2021 Jun 24;gr.268441.120.

48. Rahman R, Chirn G wei, Kanodia A, Sytnikova YA, Brembs B, Bergman CM, et al. Unique transposon landscapes are pervasive across *Drosophila melanogaster* genomes. Nucleic Acids Res. 2015 Dec 15;43(22):10655–72.

49. Mohamed M, Dang NTM, Ogyama Y, Burlet N, Mugat B, Boulesteix M, et al. A Transposon Story: From TE Content to TE Dynamic Invasion of Drosophila Genomes Using the Single-Molecule Sequencing Technology from Oxford Nanopore. Cells. 2020 Jul 25;9(8).

50. Bergman CM, Quesneville H, Anxolabéhère D, Ashburner M. Recurrent insertion and duplication generate networks of transposable element sequences in the Drosophila melanogaster genome. Genome Biol. 2006 Nov 29;7(11):R112.

51. Kaminker JS, Bergman CM, Kronmiller B, Carlson J, Svirskas R, Patel S, et al. The transposable elements of the Drosophila melanogaster euchromatin: a genomics perspective. Genome Biol. 2002;3(12):RESEARCH0084.

52. Dubnau J. The Retrotransposon storm and the dangers of a Collyer’s genome. Current Opinion in Genetics & Development. 2018 Apr;49:95–105.

53. Cardelli M. The epigenetic alterations of endogenous retroelements in aging. Mechanisms of Ageing and Development. 2018 Sep;174:30–46.

54. Gorbunova V, Seluanov A, Mita P, McKerrow W, Fenyö D, Boeke JD, et al. The role of retrotransposable elements in ageing and age-associated diseases. Nature. 2021 Aug 5;596(7870):43–53.

55. Sousa-Victor P, Ayyaz A, Hayashi R, Qi Y, Madden DT, Lunyak VV, et al. Piwi Is Required to Limit Exhaustion of Aging Somatic Stem Cells. Cell Reports. 2017 Sep;20(11):2527–37.

56. Tang X, Liu N, Qi H, Lin H. Piwi maintains homeostasis in the Drosophila adult intestine. Stem Cell Reports [Internet]. 2023 Feb 2 [cited 2023 Feb 8];0(0). Available from: https://www.cell.com/stem-cell-reports/abstract/S2213-6711(23)00004-8

57. Chung WJ, Okamura K, Martin R, Lai EC. Endogenous RNA Interference Provides a Somatic Defense against Drosophila Transposons. Current Biology. 2008 Jun;18(11):795–802.

58. Czech B, Malone CD, Zhou R, Stark A, Schlingeheyde C, Dus M, et al. An endogenous small interfering RNA pathway in Drosophila. Nature. 2008 Jun;453(7196):798–802.

59. Ghildiyal M, Seitz H, Horwich MD, Li C, Du T, Lee S, et al. Endogenous siRNAs Derived from Transposons and mRNAs in Drosophila Somatic Cells. Science. 2008 May 23;320(5879):1077–81.

60. Varoqui M, Mohamed M, Mugat B, Gourion D, Lemoine M, Pélisson A, et al. Temporal and spatial niche partitioning in a retrotransposon community of the Drosophila genome. bioRxiv. 2024 Jan 1;2024.08.14.607943.

61. Johnson WE. Origins and evolutionary consequences of ancient endogenous retroviruses. Nat Rev Microbiol. 2019 Jun;17(6):355–70.

62. Gervais L, van den Beek M, Josserand M, Sallé J, Stefanutti M, Perdigoto CN, et al. Stem Cell Proliferation Is Kept in Check by the Chromatin Regulators Kismet/CHD7/CHD8 and Trr/MLL3/4. Developmental Cell. 2019 May 20;49(4):556–573.e6.

63. Josserand M, Rubanova N, Stefanutti M, Roumeliotis S, Espenel M, Marshall OJ, et al. Chromatin state transitions in the Drosophila intestinal lineage identify principles of cell-type specification. Developmental Cell. 2023 Dec;58(24):3048–3063.e6.

64. Desset S, Meignin C, Dastugue B, Vaury C. COM, a Heterochromatic Locus Governing the Control of Independent Endogenous Retroviruses From *Drosophila melanogaster*. Genetics. 2003 Jun 1;164(2):501–9.

65. Sarot E, Payen-Groschêne G, Bucheton A, Pélisson A. Evidence for a *piwi* -Dependent RNA Silencing of the *gypsy* Endogenous Retrovirus by the *Drosophila melanogaster flamenco* Gene. Genetics. 2004 Mar 1;166(3):1313–21.

66. Senti KA, Handler D, Rafanel B, Kosiol C, Schlötterer C, Brennecke J. Functional Adaptations of Endogenous Retroviruses to the *Drosophila* Host Underlie their Evolutionary Diversification. bioRxiv. 2023 Jan 1;2023.08.03.551782.

67. The modENCODE Consortium, Roy S, Ernst J, Kharchenko PV, Kheradpour P, Negre N, et al. Identification of Functional Elements and Regulatory Circuits by Drosophila modENCODE. Science. 2010 Dec 24;330(6012):1787–97.

68. Dutta D, Dobson AJ, Houtz PL, Gläßer C, Revah J, Korzelius J, et al. Regional Cell-Specific Transcriptome Mapping Reveals Regulatory Complexity in the Adult Drosophila Midgut. Cell Reports. 2015 Jul;12(2):346–58.

69. Jiang H, Patel PH, Kohlmaier A, Grenley MO, McEwen DG, Edgar BA. Cytokine/Jak/Stat Signaling Mediates Regeneration and Homeostasis in the Drosophila Midgut. Cell. 2009 Jun;137(7):1343–55.

70. Korzelius J, Naumann SK, Loza-Coll MA, Chan JS, Dutta D, Oberheim J, et al. *Escargot* maintains stemness and suppresses differentiation in *Drosophila* intestinal stem cells. EMBO J. 2014 Dec 17;33(24):2967–82.

71. Antonello ZA, Reiff T, Ballesta-Illan E, Dominguez M. Robust intestinal homeostasis relies on cellular plasticity in enteroblasts mediated by miR-8-Escargot switch. The EMBO Journal. 2015 Aug 4;34(15):2025–41.

72. Li W, Prazak L, Chatterjee N, Grüninger S, Krug L, Theodorou D, et al. Activation of transposable elements during aging and neuronal decline in Drosophila. Nat Neurosci. 2013 May;16(5):529–31.

73. Treiber CD, Waddell S. Resolving the prevalence of somatic transposition in Drosophila. eLife. 2017 Jul 25;6:e28297.

74. Jones BC, Wood JG, Chang C, Tam AD, Franklin MJ, Siegel ER, et al. A somatic piRNA pathway in the Drosophila fat body ensures metabolic homeostasis and normal lifespan. Nat Commun. 2016 Dec;7(1):13856.

75. Chang YH, Keegan RM, Prazak L, Dubnau J. Cellular labeling of endogenous retrovirus replication (CLEVR) reveals de novo insertions of the gypsy retrotransposable element in cell culture and in both neurons and glial cells of aging fruit flies. PLOS Biology. 2019 May 16;17(5):e3000278.

76. Yang N, Srivastav SP, Rahman R, Ma Q, Dayama G, Li S, et al. Transposable element landscapes in aging Drosophila. PLOS Genetics. 2022 Mar 3;18(3):e1010024.

77. Rigal J, Martin Anduaga A, Bitman E, Rivellese E, Kadener S, Marr MT. Artificially stimulating retrotransposon activity increases mortality and accelerates a subset of aging phenotypes in Drosophila. Botchan MR, Isales C, editors. eLife. 2022 Aug 18;11:e80169.

78. Schneider BK, Sun S, Lee M, Li W, Skvir N, Neretti N, et al. Expression of retrotransposons contributes to aging in Drosophila. Genetics. 2023 Jun 1;224(2):iyad073.

79. Schauer SN, Carreira PE, Shukla R, Gerhardt DJ, Gerdes P, Sanchez-Luque FJ, et al. L1 retrotransposition is a common feature of mammalian hepatocarcinogenesis. Genome Res. 2018 May;28(5):639–53.

80. Lanciano S, Philippe C, Sarkar A, Pratella D, Domrane C, Doucet AJ, et al. Locus-level L1 DNA methylation profiling reveals the epigenetic and transcriptional interplay between L1s and their integration sites. Cell Genomics. 2024 Feb;100498.

81. Ewing AD, Smits N, Sanchez-Luque FJ, Faivre J, Brennan PM, Richardson SR, et al. Nanopore Sequencing Enables Comprehensive Transposable Element Epigenomic Profiling. Molecular Cell. 2020 Dec 3;80(5):915–928.e5.

82. Glaser J, Cova G, Fauler B, Prada-Medina CA, Stanislas V, Phan MHQ, et al. Enhancer adoption by an LTR retrotransposon generates viral-like particles causing developmental limb phenotypes. bioRxiv. 2024 Jan 1;2024.09.13.612906.

83. Sun X, Wang X, Tang Z, Grivainis M, Kahler D, Yun C, et al. Transcription factor profiling reveals molecular choreography and key regulators of human retrotransposon expression. Proc Natl Acad Sci USA [Internet]. 2018 Jun 12 [cited 2024 Aug 25];115(24). Available from: https://pnas.org/doi/full/10.1073/pnas.1722565115

84. Hermant C, Torres-Padilla ME. TFs for TEs: the transcription factor repertoire of mammalian transposable elements. Genes Dev. 2021 Jan 1;35(1–2):22–39.

85. Becker KG, Swergold G, Ozato K, Thayer RE. Binding of the ubiquitous nuclear transcription factor YY1 to a *cis* regulatory sequence in the human LINE-1 transposable element. Hum Mol Genet. 1993;2(10):1697–702.

86. Yang N. An important role for RUNX3 in human L1 transcription and retrotransposition. Nucleic Acids Research. 2003 Aug 15;31(16):4929–40.

87. Wylie A, Jones AE, D’Brot A, Lu WJ, Kurtz P, Moran JV, et al. p53 genes function to restrain mobile elements. Genes Dev. 2016 Jan 1;30(1):64–77.

88. Tchenio T. Members of the SRY family regulate the human LINE retrotransposons. Nucleic Acids Research. 2000 Jan 15;28(2):411–5.

89. Muotri AR, Marchetto MCN, Coufal NG, Oefner R, Yeo G, Nakashima K, et al. L1 retrotransposition in neurons is modulated by MeCP2. Nature. 2010 Nov;468(7322):443–6.

90. Kuwabara T, Hsieh J, Muotri A, Yeo G, Warashina M, Lie DC, et al. Wnt-mediated activation of NeuroD1 and retro-elements during adult neurogenesis. Nat Neurosci. 2009 Sep;12(9):1097–105.

91. Tang H, Yang J, Xu J, Zhang W, Geng A, Jiang Y, et al. The transcription factor PAX5 activates human LINE1 retrotransposons to induce cellular senescence. EMBO Rep. 2024 Jun 12;25(8):3263– 75.

92. Micchelli CA, Perrimon N. Evidence that stem cells reside in the adult Drosophila midgut epithelium. Nature. 2006 Jan;439(7075):475–9.

93. Ohlstein B, Spradling A. The adult Drosophila posterior midgut is maintained by pluripotent stem cells. Nature. 2006 Jan;439(7075):470–4.

94. Kosman D, Ip YT, Levine M, Arora K. Establishment of the Mesoderm-Neuroectoderm Boundary in the Drosophila Embryo. Science. 1991 Oct 4;254(5028):118–22.

95. Rembold M, Ciglar L, Yáñez-Cuna JO, Zinzen RP, Girardot C, Jain A, et al. A conserved role for Snail as a potentiator of active transcription. Genes Dev. 2014 Jan 15;28(2):167–81.

96. Rech GE, Radío S, Guirao-Rico S, Aguilera L, Horvath V, Green L, et al. Population-scale long-read sequencing uncovers transposable elements associated with gene expression variation and adaptive signatures in Drosophila. Nat Commun. 2022 Apr 12;13(1):1948.

97. Stewart C, Kural D, Strömberg MP, Walker JA, Konkel MK, Stütz AM, et al. A Comprehensive Map of Mobile Element Insertion Polymorphisms in Humans. Malik HS, editor. PLoS Genet. 2011 Aug 18;7(8):e1002236.

98. Rishishwar L, Tellez Villa CE, Jordan IK. Transposable element polymorphisms recapitulate human evolution. Mobile DNA. 2015 Dec;6(1):21.

99. Sudmant PH, Rausch T, Gardner EJ, Handsaker RE, Abyzov A, Huddleston J, et al. An integrated map of structural variation in 2,504 human genomes. Nature. 2015 Oct 1;526(7571):75–81.

100. Chu C, Borges-Monroy R, Viswanadham VV, Lee S, Li H, Lee EA, et al. Comprehensive identification of transposable element insertions using multiple sequencing technologies. Nat Commun. 2021 Jun 22;12(1):3836.

101. Zhou X, Sam TW, Lee AY, Leung D. Mouse strain-specific polymorphic provirus functions as cis-regulatory element leading to epigenomic and transcriptomic variations. Nat Commun. 2021 Nov 9;12(1):6462.

102. Alonge M, Wang X, Benoit M, Soyk S, Pereira L, Zhang L, et al. Major Impacts of Widespread Structural Variation on Gene Expression and Crop Improvement in Tomato. Cell. 2020 Jul;182(1):145–161.e23.

103. Domínguez M, Dugas E, Benchouaia M, Leduque B, Jiménez-Gómez JM, Colot V, et al. The impact of transposable elements on tomato diversity. Nat Commun. 2020 Aug 13;11(1):4058.

104. Burns KH. Our Conflict with Transposable Elements and Its Implications for Human Disease. Annu Rev Pathol Mech Dis. 2020 Jan 24;15(1):51–70.

105. Bischof J, Maeda RK, Hediger M, Karch F, Basler K. An optimized transgenesis system for *Drosophila* using germ-line-specific φC31 integrases. Proc Natl Acad Sci USA. 2007 Feb 27;104(9):3312–7.

106. Schindelin J, Arganda-Carreras I, Frise E, Kaynig V, Longair M, Pietzsch T, et al. Fiji: an open-source platform for biological-image analysis. Nat Methods. 2012 Jul;9(7):676–82.

107. Buchon N, Osman D, David FPA, Yu Fang H, Boquete JP, Deplancke B, et al. Morphological and Molecular Characterization of Adult Midgut Compartmentalization in Drosophila. Cell Reports. 2013 May;3(5):1725–38.

108. De Coster W, D’Hert S, Schultz DT, Cruts M, Van Broeckhoven C. NanoPack: visualizing and processing long-read sequencing data. Berger B, editor. Bioinformatics. 2018 Aug 1;34(15):2666–9.

109. Öztürk-Çolak A, Marygold SJ, Antonazzo G, Attrill H, Goutte-Gattat D, Jenkins VK, et al. FlyBase: updates to the *Drosophila* genes and genomes database. Wood V, editor. GENETICS. 2024 May 7;227(1):iyad211.

110. Li H. Minimap2: pairwise alignment for nucleotide sequences. Birol I, editor. Bioinformatics. 2018 Sep 15;34(18):3094–100.

111. Danecek P, Bonfield JK, Liddle J, Marshall J, Ohan V, Pollard MO, et al. Twelve years of SAMtools and BCFtools. GigaScience. 2021 Jan 29;10(2):giab008.

112. De Coster W, Rademakers R. NanoPack2: population-scale evaluation of long-read sequencing data. Alkan C, editor. Bioinformatics. 2023 May 4;39(5):btad311.

113. Leger A, Leonardi T. pycoQC, interactive quality control for Oxford Nanopore Sequencing. JOSS. 2019 Feb 28;4(34):1236.

114. Storer J, Hubley R, Rosen J, Wheeler TJ, Smit AF. The Dfam community resource of transposable element families, sequence models, and genome annotations. Mobile DNA. 2021 Jan 12;12(1):2.

115. Bailly-Bechet M, Haudry A, Lerat E. “One code to find them all”: a perl tool to conveniently parse RepeatMasker output files. Mobile DNA. 2014 Dec;5(1):13.

116. Han S, Basting PJ, Dias GB, Luhur A, Zelhof AC, Bergman CM. Transposable element profiles reveal cell line identity and loss of heterozygosity in Drosophila cell culture. Genetics. 2021 Oct 1;219(2):iyab113.

117. Rozewicki J, Li S, Amada KM, Standley DM, Katoh K. MAFFT-DASH: integrated protein sequence and structural alignment. Nucleic Acids Research. 2019 May 7;gkz342.

118. Rice P, Longden I, Bleasby A. EMBOSS: the European Molecular Biology Open Software Suite. Trends Genet. 2000 Jun;16(6):276–7.

119. Robinson JT. Integrative genomics viewer. 2011;

120. Darling ACE, Mau B, Blattner FR, Perna NT. Mauve: Multiple Alignment of Conserved Genomic Sequence With Rearrangements. Genome Res. 2004 Jul;14(7):1394–403.

121. Dobin A, Davis CA, Schlesinger F, Drenkow J, Zaleski C, Jha S, et al. STAR: ultrafast universal RNA-seq aligner. Bioinformatics. 2013 Jan 1;29(1):15–21.

122. Robinson MD, McCarthy DJ, Smyth GK. edgeR : a Bioconductor package for differential expression analysis of digital gene expression data. Bioinformatics. 2010 Jan 1;26(1):139–40.

123. Love MI, Huber W, Anders S. Moderated estimation of fold change and dispersion for RNA-seq data with DESeq2. Genome Biology. 2014 Dec 5;15(12):550.

124. Liao Y, Smyth GK, Shi W. featureCounts: an efficient general purpose program for assigning sequence reads to genomic features. Bioinformatics. 2014 Apr 1;30(7):923–30.

125. Gershenzon NI, Trifonov EN, Ioshikhes IP. The features of Drosophila core promoters revealed by statistical analysis. BMC Genomics. 2006 Dec;7(1):161.

126. Tan G, Lenhard B. TFBSTools: an R/bioconductor package for transcription factor binding site analysis. Bioinformatics. 2016 May 15;32(10):1555–6.

127. Rauluseviciute I, Riudavets-Puig R, Blanc-Mathieu R, Castro-Mondragon JA, Ferenc K, Kumar V, et al. JASPAR 2024: 20th anniversary of the open-access database of transcription factor binding profiles. Nucleic Acids Research. 2024 Jan 5;52(D1):D174–82.

128. Ramírez F, Ryan DP, Grüning B, Bhardwaj V, Kilpert F, Richter AS, et al. deepTools2: a next generation web server for deep-sequencing data analysis. Nucleic Acids Res. 2016 Jul 8;44(W1):W160–5.

