## Supplementary material for "An endogenous retroviral element co-opts an upstream regulatory sequence to achieve somatic expression and mobility": Rubanova etal-Supplementary File

bioRxiv

### Supplementary File:

Supplementary Figures 1-11
Supplementary information: Full *Drosophila* genotypes used in this study

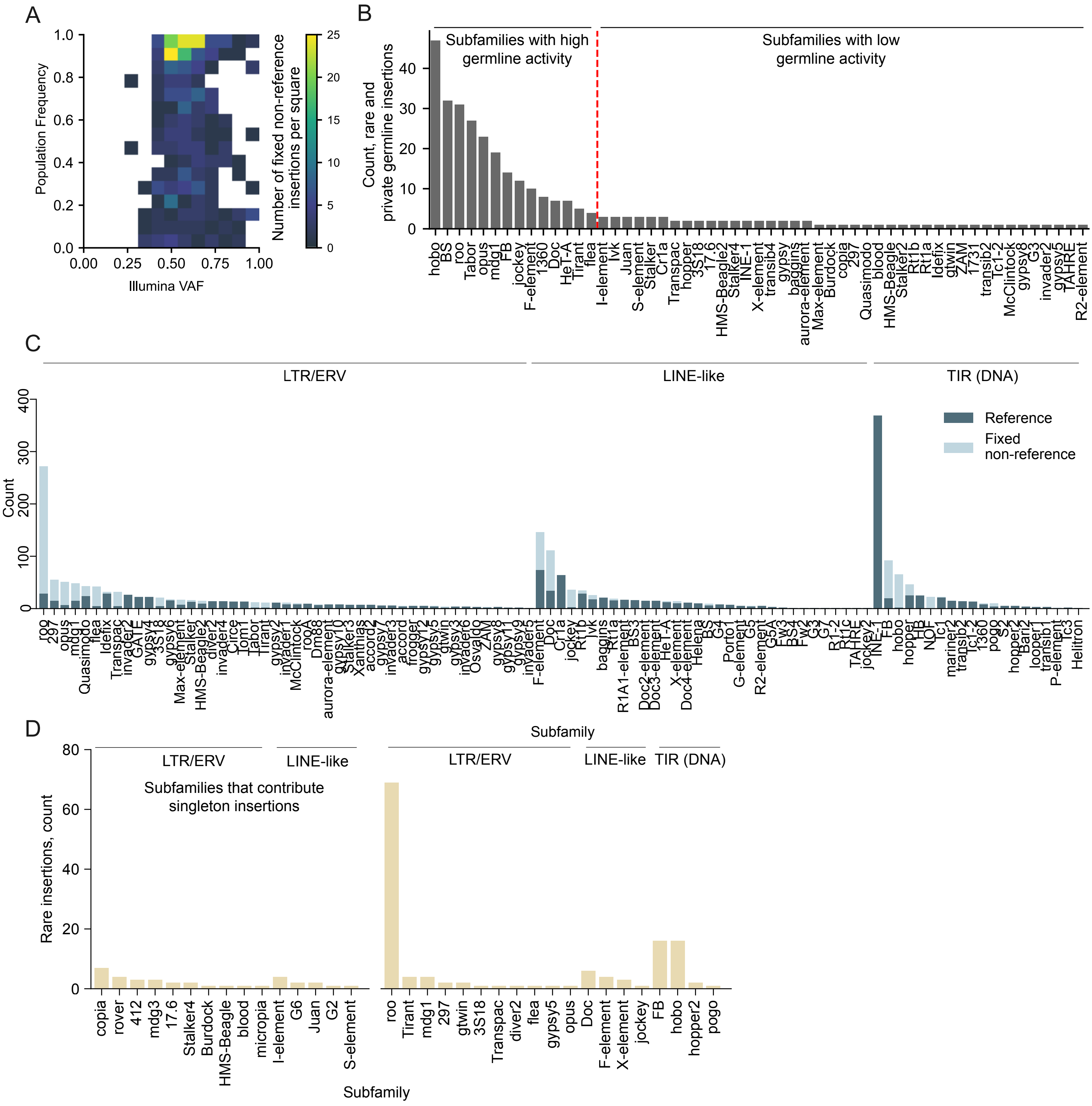


**Supplementary Figure 1. The landscape of the full-length TE insertions in the *ProsGFP* strain.**

(A) Density plot for 747 out of 1227 of fixed non-reference insertions detected in our previously published Illumina DNA sequencing libraries, representing Illumina VAF and population frequency. Each square represents an interval of Illumina VAF and population frequency, with the color code indicatingthe number of fixed non-reference insertions within the corresponding interval.

(B) The number of rare germline and private germline insertions per sub-family detected in the Illumina DNA sequencing libraries. Private germline insertions are defined as those detected exactly in one sample pair (gut and head sampes from one fly). Rare germline insertions are those detected in more than 1 but fewer than 3 sample pairs (see Materials and Methods section).

(C) The number of detected fixed full-length reference (dark blue) and fixed non-reference (light blue) insertions of TE sub-families that did not contribute somatic insertions.

(D) The number of detected rare insertions per sub-family. Sub-families are grouped into those contributing singleton insertions (left side) and all other sub-families (right side).

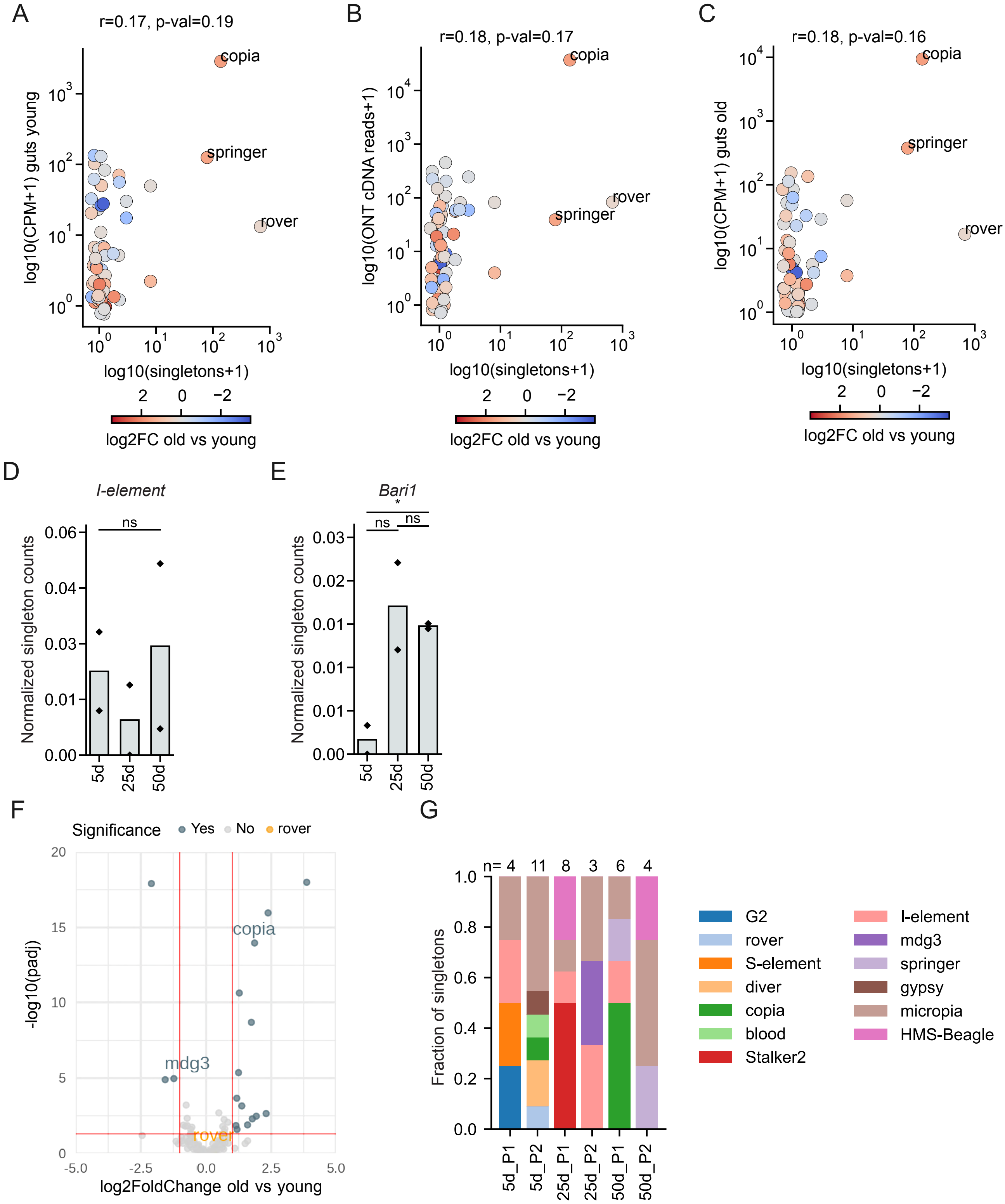


**Supplementary Figure 2. TE transcriptional activity and mobility in tissues of different age groups.**

(A-C) Correlation between the total number of singleton insertions and the expression level of TE sub-families in each of the three age groups. The color code represents the log2 fold change of the expression level in old vs young flies. (A) Young flies. (B) Mid-aged flies. (C) Old flies.

(D-E) Normalized singleton counts in the three age groups. The bar represents the average between two replicates. (D) LINE-like *I-element*. (E) DNA element *Bari1*.

(F) Volcano plot representing differential expression of TE sub-families in old *vs* young head tissue of the *ProsGFP* strain. Significantly up- and downregulated sub-families are indicated in blue, and sub-families which were detected as mobile in the head tissue are labeled.

(G) Fraction of singleton insertions for each sub-family detected in head samples per age group per replicate. P1 and P2 are two biological replicates (pool 1 and 2). n= the total number of singletons in each pool.

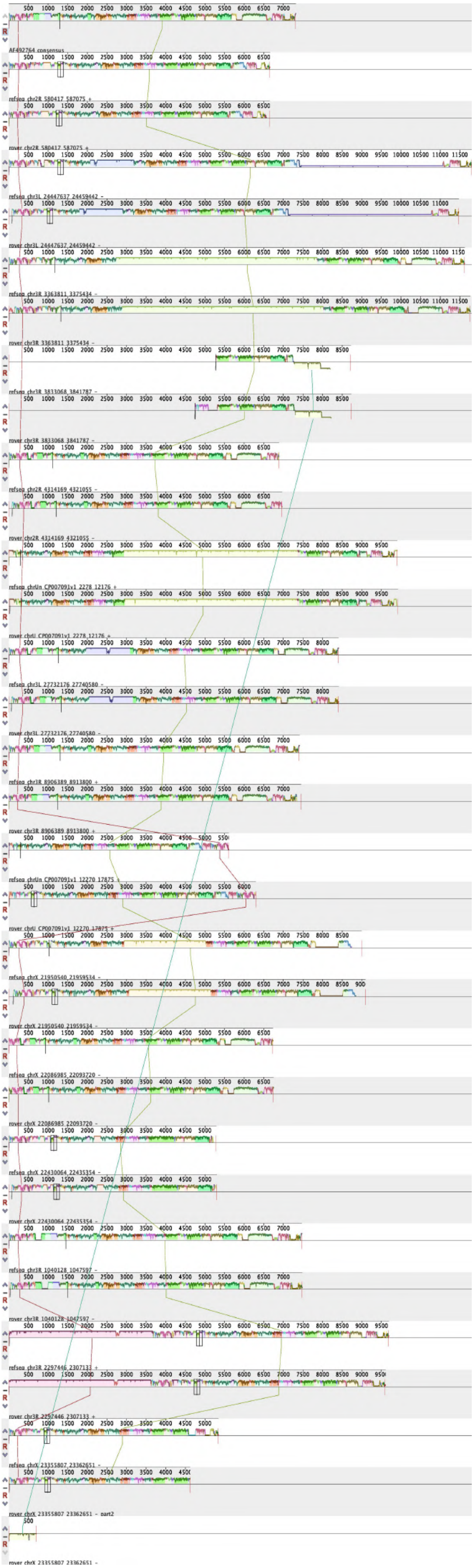

**Supplementary Figure 3. Multiple sequence alignment shows structural variation in the reference *rover* insertions present in the *ProsGFP* genome.**

The MAUVE graphical visualization of the multiple sequence alignment of the *rover* consensus sequence and the fixed reference insertions. The first row represents the *rover* consensus sequence. Subsequent rows are paired for each reference insertion: the first row in each pair represents the sequence extracted from the reference genome, and the second row represents the sequence assembled from the ONT reads. Colors indicate regions with similar sequence.

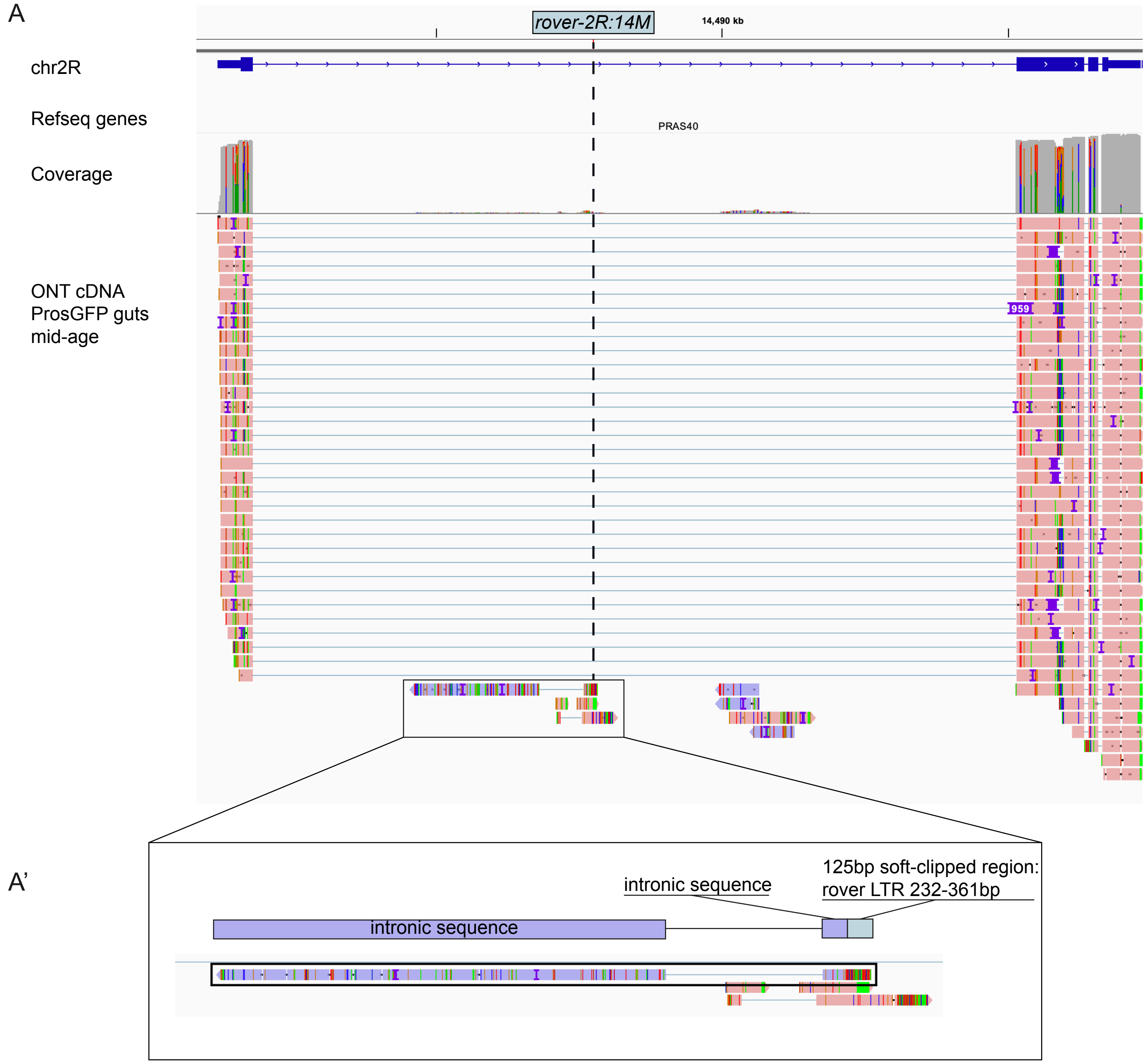


**Supplementary Figure 4. Visualisation of the ONT cDNA reads from gut libraries of the *ProsGFP* background covering *PRAS40* locus.**

(A) IGV visualization of the aligned ONT cDNA reads showing unperturbed transcription and splicing of the *PRAS40* locus in the presence of the *rover-2R-14M* insertion. Position of the *rover-2R-14M* insertion within the first intron is indicated with the dashed line.

(A') Highlight of a single read (black box) that starts in the *rover-2R-14M* LTR region (125bp soft-clipped region) and continues into the *PRAS40* intron, suggesting that *rover-2R-14M* can potentially provide an aberrant transcription start site. The remaining two reads visible in the panel do not align to *rover-2R-14M*.

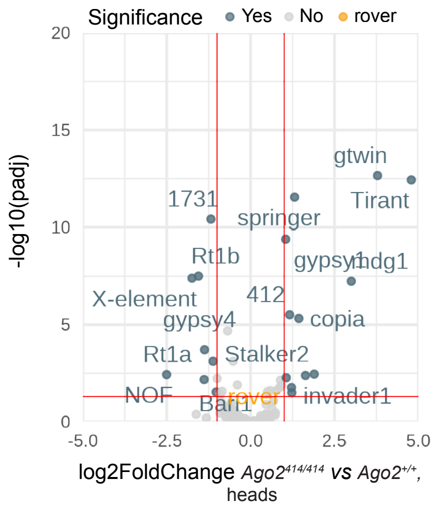


**Supplementary Figure 5. Differential TE expression in *Ago2* mutant condition.**

Volcano plot representing differential expression of TE sub-families in the head tissue in the *Ago2* mutant condition (*rover-2R:14M*$^{+/+}$*;Ago2*$^{414/414}$ *vs rover-2R:14M*$^{+/+}$*;Ago2*$^{+/+}$). Both genotypes carry the *rover-2R:14M* active copy. Significantly up- or down-regulated TE sub-families are indicated in blue and labeled.

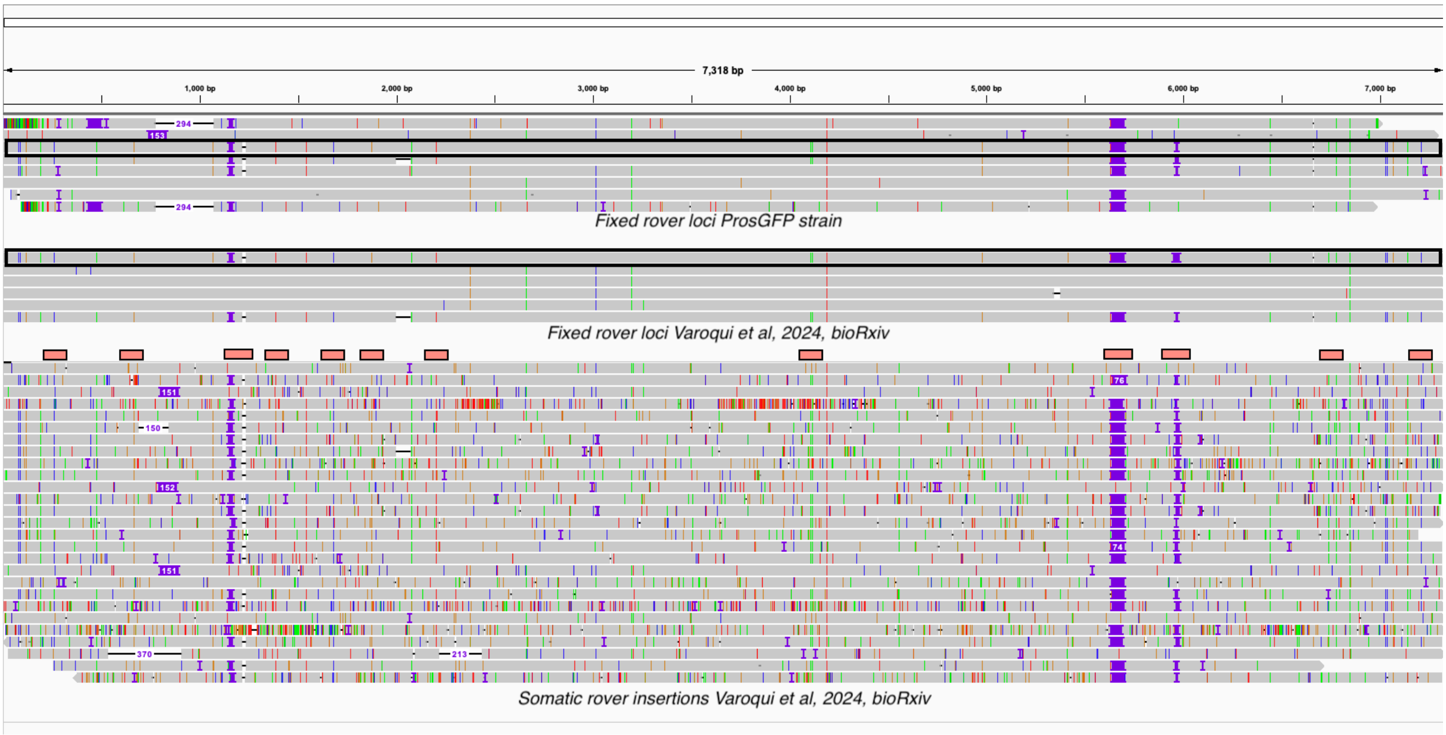


**Supplementary Figure 6. ONT DNA-seq evidence that *rover-2R-14M* may potentially be somatically active in the genetic background used in Varoqui et al. 2024.**

IGV visualization of alignments of *rover* fixed and putative somatic insertions from Varoqui et al. 2024, bioRxiv (doi: https://doi.org/10.1101/2024.08.14.607943), as well as fixed *rover* insertions present in the *ProsGFP* strain (this study). *Rover-2R-14M* copies are highlighted with black border around them. The positions of the main sequence variants that distinguish *rover-2R-14M* are highlighted with pink bars.

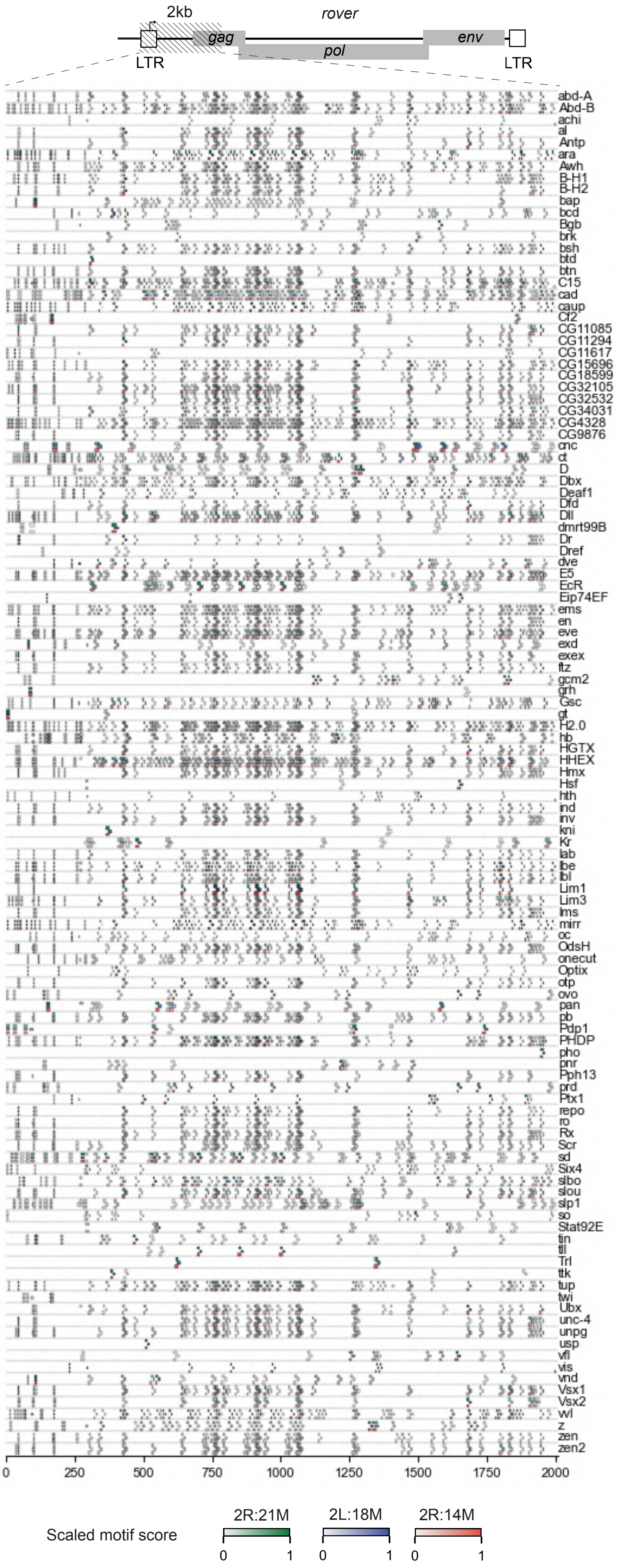


**Supplementary Figure 7. Motif analysis of the 2kb-internal region of the fixed *rover* loci present in the *ProsGFP* strain.**

Putative transcription factor binding sites of the transcription factors with expression levels lower than 10 RPKM in at least one gut cell type. 2kb region of the internal sequence of three *rover* loci was analysed: *2R-14M* (red), *2L-18M* (blue), *2R-21M* (green).

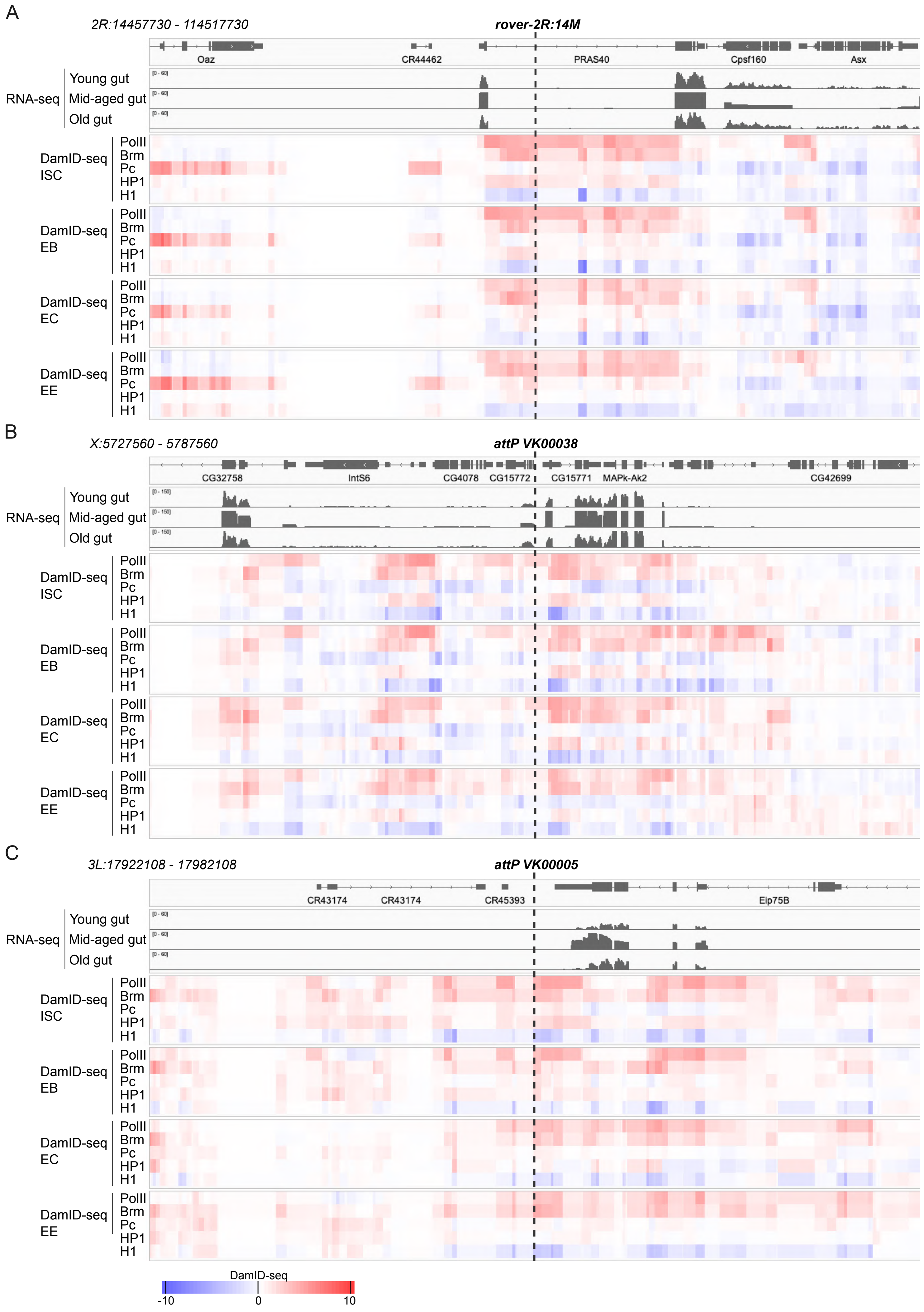
A
2R:14457730 - 114517730
rover-2R:14M
Oaz
CR44462
PRAS40
Cpsf160
Asx
RNA-seq
Young gut
Mid-aged gut
Old gut
DamID-seq ISC
DamID-seq EB
DamID-seq EC
DamID-seq EE
PolII
Brm
Pc
HP1
H1
B
X:5727560 - 5787560
attP VK00038
CG32758
IntS6
CG4078
CG15772
CG15771
MAPk-Ak2
CG42699
C
3L:17922108 - 17982108
attP VK00005
CR43174
CR43174
CR45393
Eip75B
DamID-seq
-10
0
10

**Supplementary Figure 8. Chromatin landscapes at the *rover2R:14M* insertion site (A), and the two attP landing sites in which *rover-lacZ* reporters were inserted on the X (B) and the 3rd chromosome (C).**

For each genomic site, a region of 60 kb is illustrated, with the insertion site in the middle (dotted line). Each panel shows (from top to bottom): annotated genes (dm6 reference genome); RNA-seq tracks from young (Illumina), mid-aged (ONT cDNA) and old (Illumina) guts; and DamID-seq binding profiles of Polymerase II (PolII), Brahma (Brm), Polycomb (Pc), Heterochromatin protein 1a (HP1) and histone H1 (H1), for different gut cell types. ISC, intestinal stem cells; EB, enteroblasts; EC, enterocytes; EE, eneteroendocrine cells. DamID-seq data are from Josserand et al, 2023 (doi: 10.1016/j.devcel.2023.11.005).

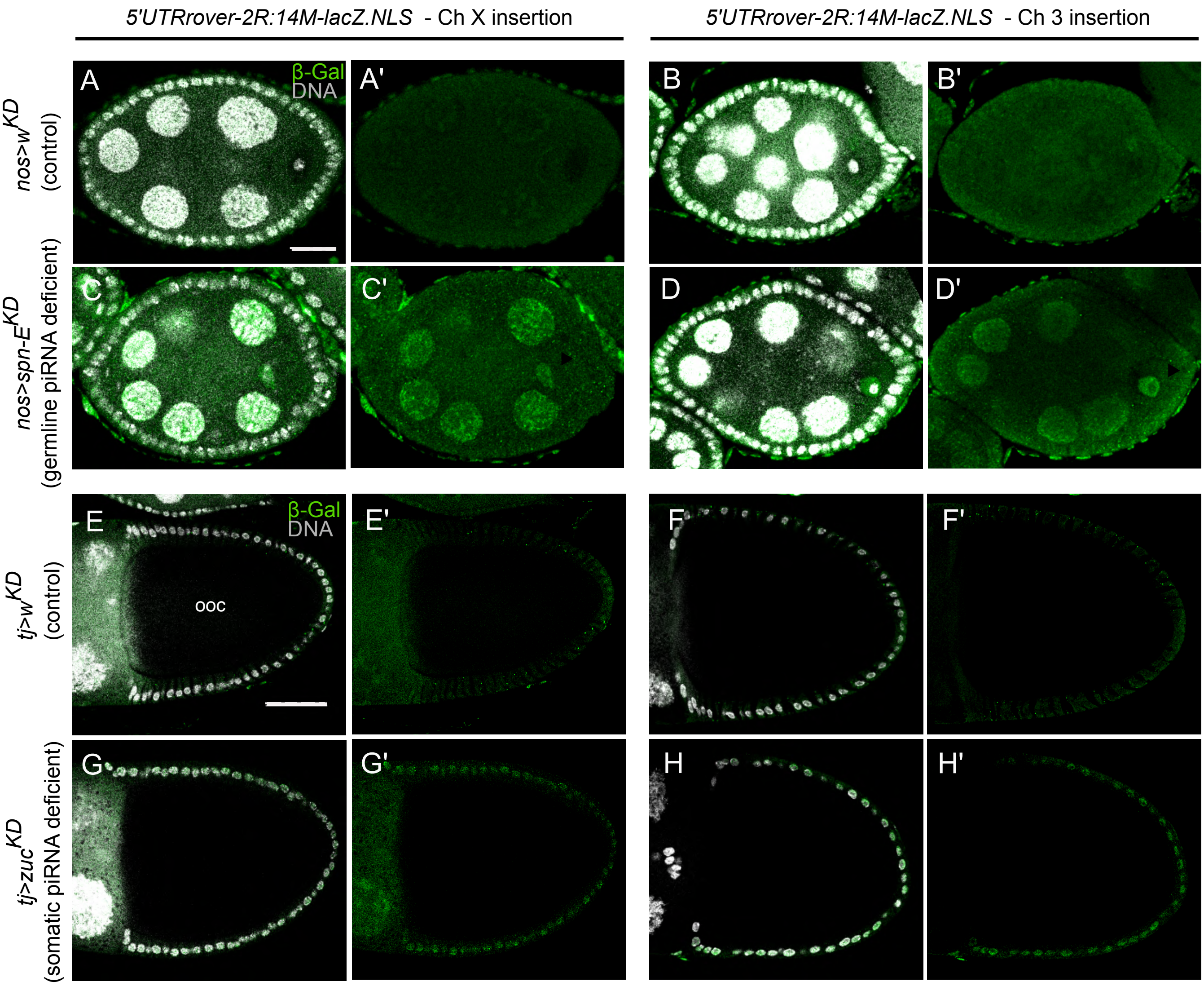


**Supplementary Figure 9.** ***rover-2R:14M-lacZ.NLS*** **reporter is expressed in ovaries with non-functional piRNA pathway.**

Whole mount immunostaining of adult ovaries (one confocal section) from females carrying the *rover-2R:14M-lacZ.NLS* construct inserted on the X or on the 3rd chromosome. β-galactosidase (green), DAPI (grey). Anterior is left. Scale bar: 50 μm.

(A-D') *rover-2R:14M-lacZ.NLS* reporter expression in stage 6/7 egg chambers in ovaries with a germline knock-down of *white* (control, A-B') or *spnE* (piRNA pathway component, C-D') with the germline *nosGAL4* driver. Reporter expression is detected in germ cell nuclei (including the oocyte, arrow) only in the piRNA pathway-deficient genetic context, for both chromosomal reporter insertions. Note that, in the oocyte, the anti-β-galactosidase staining is larger than the DAPI signal, since only part of oocyte nucleus is stained by DAPI in ovaries. Scale bar: 25 μm.

(E-H') *rover-2R:14M-lacZ.NLS* reporter expression in stage 9/10 egg chamber in ovaries with somatic knock-down of *white* (control, E-F') or *zuc* (piRNA pathway component, G-H') with the *tjGAL4* driver. Anti-β-galactosidase staining is detected in follicle somatic cells surrounding the oocyte (ooc) in the piRNA pathway-deficient genetic context, for both chromosomal reporter insertions.

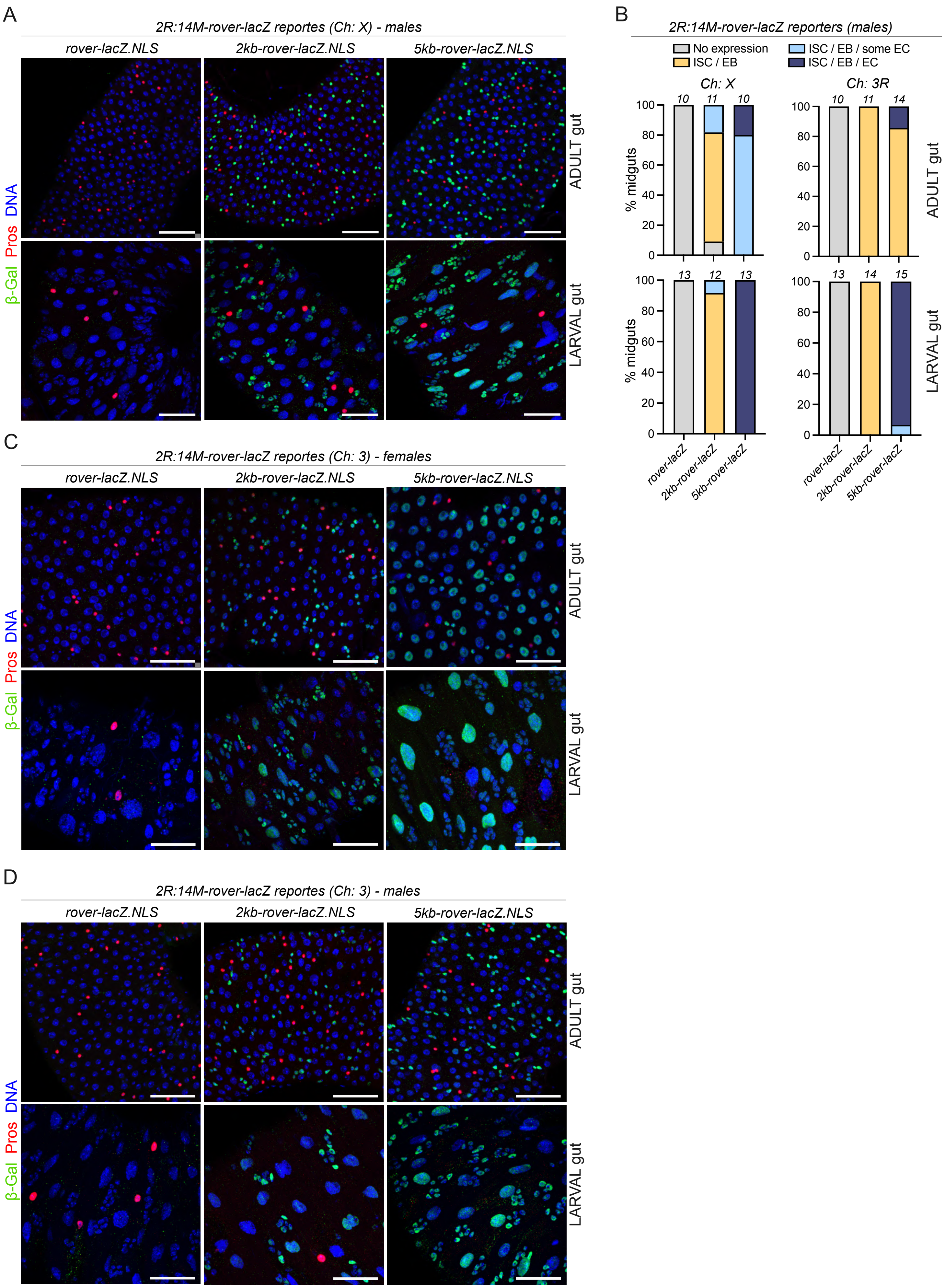
A
2R:14M-rover-lacZ reportes (Ch: X) - males
rover-lacZ.NLS
2kb-rover-lacZ.NLS
5kb-rover-lacZ.NLS
ADULT gut
LARVAL gut
β-Gal Pros DNA
B
2R:14M-rover-lacZ reporters (males)
No expression
ISC / EB
ISC / EB / some EC
ISC / EB / EC
Ch: X
Ch: 3R
% midguts
ADULT gut
LARVAL gut
10 11 10
10 11 14
13 12 13
13 14 15
rover-lacZ
2kb-rover-lacZ
5kb-rover-lacZ
C
2R:14M-rover-lacZ reportes (Ch: 3) - females
rover-lacZ.NLS
2kb-rover-lacZ.NLS
5kb-rover-lacZ.NLS
ADULT gut
LARVAL gut
β-Gal Pros DNA
D
2R:14M-rover-lacZ reportes (Ch: 3) - males
rover-lacZ.NLS
2kb-rover-lacZ.NLS
5kb-rover-lacZ.NLS
ADULT gut
LARVAL gut
β-Gal Pros DNA

**Supplementary Figure 10. *rover-2R:14M-lacZ.NLS* reporter expression in the fly larval and adult midguts.**

Tissues were stained for β-galactosidase (β-gal, green), Prospero (Pros, red) and DNA (blue). Scale bar: 50 μm.

(A) Representative images of male adult and larval midguts carrying *rover-2R:14M-lacZ* reporter constructs inserted on the X chromosome.

(B) Quantifications of the expression patterns of *rover-2R:14M-lacZ.NLS* reporter constructs in male adult and larval midguts. Numbers of midguts scored are indicated above each bar.

(C-D) Representative images of female (C) and male (D) adult and larval midguts carrying *rover-2R:14M-lacZ* reporter constructs inserted on the 3rd chromosome.

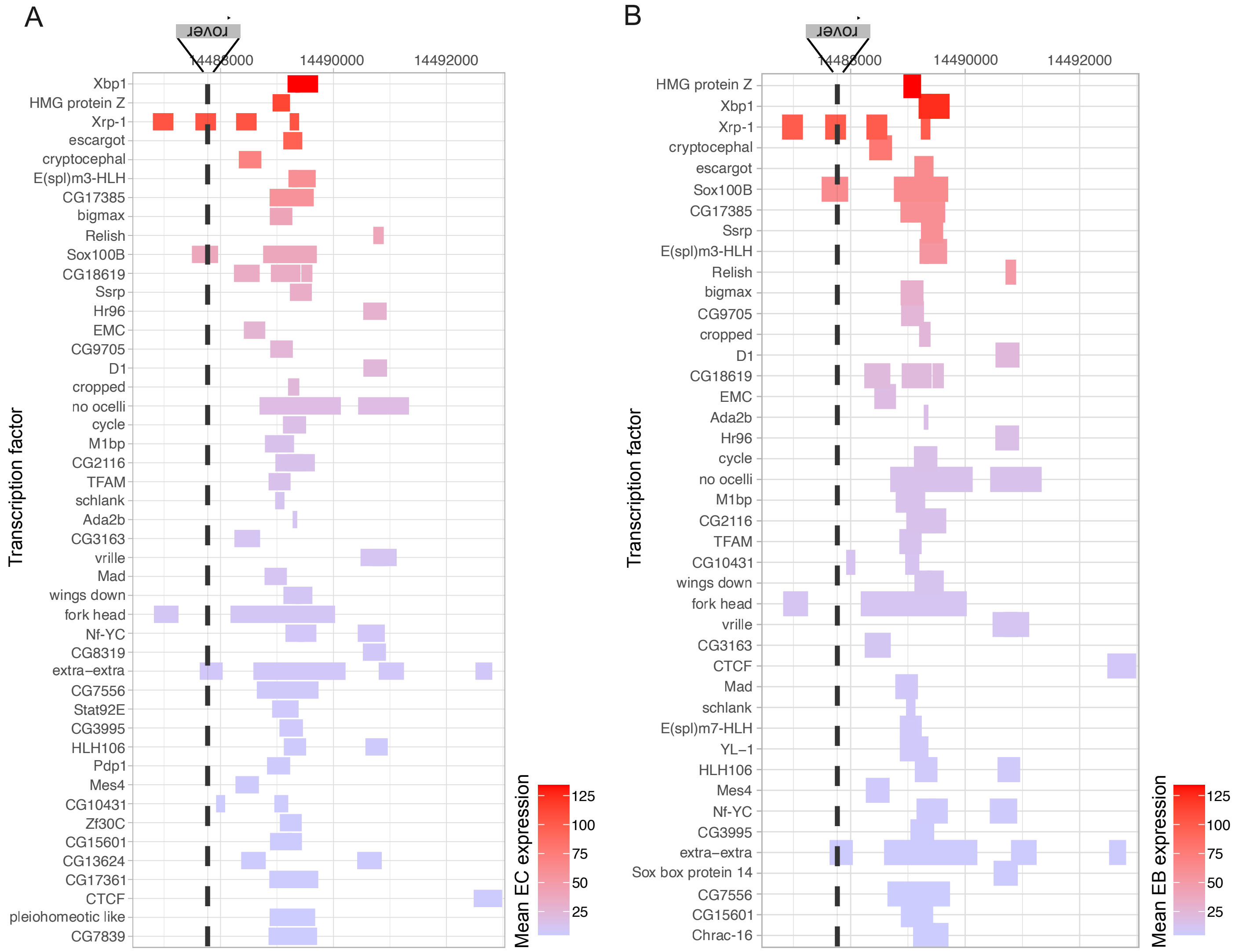


**Supplementary Figure 11. Transcription factor binding sites in the genomic region upstream of the *rover-2R:14M* locus.**

Published ChiP-seq datasets (modENCODE, DOI: 10.1126/science.1198374) were analyzed as for Figure 6A. Bars indicating TF binding sites were ordered and colored based on mean transcription factor expression levels in differentiated enterocytes (EC) (A) or EC progenitors, enteroblasts (EB) (B). Transcriptomic data were from Dutta et al, 2015 (doi: 10.1016/j.celrep.2015.06.009).

### Supplementary information

Full *Drosophila* genotypes used in this study as represented in figure panels

Figure 1:

*$w^{1118}$; rover2R:14M /UAS-2xGFP/+ ; ProsGAL4/+* (all panels)

Figure 2:

*$w^{1118}$; rover2R:14M /UAS-2xGFP/+ ; ProsGAL4/+* (all panels)

Figure 3:

A-C and F-I:

*$w^{1118}$; rover2R:14M /UAS-2xGFP/+ ; ProsGAL4/+*

D and E:

*$w^{1118}$; rover2R:14M /UAS-2xGFP/+ ; ProsGAL4/+*

*$w^{1118}$*

J-K:

*$w^{1118}$; rover2R:14M /rover2R:14M;*

*$w^{1118}$; rover2R:14M /rover2R:14M; $Ago2^{414}$ /$Ago2^{414}$*

Figure 5:

C and F:

*5'UTRrover-LacZ.NLS /+ ; ;*

D and G:

*2kb-5'UTRrover-LacZ.NLS /+ ; ;*

E and H:

*5kb-5'UTRrover-LacZ.NLS /+ ; ;*

Figure 6:

B-B':

*2kb-5'UTRrover-LacZ.NLS /+ ; EsgGAL4 /+ ; UAS-white RNAi /TubGAL80ts UAS-GFP*

C-C':

*2kb-5'UTRrover-LacZ.NLS /+ ; EsgGAL4 /+ ; UAS-esg RNAi /TubGAL80ts UAS-GFP*

D-D':

*5kb-5'UTRrover-LacZ.NLS /+ ; EsgGAL4 /+ ; UAS-white RNAi /TubGAL80ts UAS-GFP*

E-E':

*5kb-5'UTRrover-LacZ.NLS /+ ; EsgGAL4 /+ ; UAS-esg RNAi /TubGAL80ts UAS-GFP*

Supplementary Figure 1:

*$w^{1118}$; rover2R:14M /UAS-2xGFP/+ ; ProsGAL4/+* (all panels)

Supplementary Figure 2:

*$w^{1118}$; rover2R:14M /UAS-2xGFP/+ ; ProsGAL4/+* (all panels)

Supplementary Figure 4:

*$w^{1118}$; rover2R:14M /UAS-2xGFP/+ ; ProsGAL4/+* (all panels)

Supplementary Figure 5:

*$w^{1118}$; rover2R:14M /rover2R:14M;*

*$w^{1118}$; rover2R:14M /rover2R:14M; $Ago2^{414}$ /$Ago2^{414}$*

Supplementary Figure 9:

A-A':

*5'UTRrover-LacZ.NLS /+ ; nanosGAL4 /+ ; UAS-white RNAi /+*

B-B':
; *nanosGAL4 /+ ; UAS-white RNAi / 5'UTRrover-LacZ.NLS*
C-C':
*5'UTRrover-LacZ.NLS /+ ; nanosGAL4 /+ ; UAS-spn-E RNAi /+*
D-D':
; *nanosGAL4 /+ ; UAS-spn-E RNAi / 5'UTRrover-LacZ.NLS*
E-E':
*5'UTRrover-LacZ.NLS /+ ; tjGAL4 /+ ; UAS-white RNAi/+*
F-F':
; *nanosGAL4 /+ ; UAS-white RNAi / 5'UTRrover-LacZ.NLS*
G-G':
*5'UTRrover-LacZ.NLS /+ ; tjGAL4 /+ ; UAS-zuc RNAi/+*
H-H':
; *nanosGAL4 /+ ; UAS-zuc RNAi / 5'UTRrover-LacZ.NLS*

Supplementary Figure 10:
A:
*5'UTRrover-LacZ.NLS /Y ; ;*
*2kb-5'UTRrover-LacZ.NLS /Y ; ;*
*5kb-5'UTRrover-LacZ.NLS /Y ; ;*
C and D:
*; ; 5'UTRrover-LacZ.NLS / 5'UTRrover-LacZ.NLS*
*; ; 2kb-5'UTRrover-LacZ.NLS / 2kb-5'UTRrover-LacZ.NLS*
*; ; 5kb-5'UTRrover-LacZ.NLS / 5kb-5'UTRrover-LacZ.NLS*
